# Apremilast prevents blistering in human epidermis by stabilization of keratinocyte adhesion in pemphigus

**DOI:** 10.1101/2022.02.07.478931

**Authors:** Anna M. Sigmund, Markus Winkler, Sophia Engelmayer, Desalegn T. Egu, Daniela Kugelmann, Mariya Y. Radeva, Franziska C. Bayerbach, Stefan Kotschi, Sunil Yeruva, Enno Schmidt, Amir S. Yazdi, Kamran Ghoreschi, Franziska Vielmuth, Jens Waschke

## Abstract

Pemphigus vulgaris (PV) is a life-threatening blistering skin disease caused by autoantibodies (PV-IgG) destabilizing desmosomal adhesion. Current therapies focus on suppression of autoantibody formation and thus treatments directly stabilizing keratinocyte adhesion would fulfill an unmet medical need. We here demonstrate that apremilast, a phosphodiesterase 4 inhibitor used e.g. in psoriasis, prevents blistering in PV. Apremilast abrogated PV-IgG-induced loss of keratinocyte cohesion in *ex-vivo* epidermis and *in vitro*. This was paralleled by inhibition of keratin retraction and desmosome splitting but affected neither desmoglein (Dsg) depletion nor Dsg3 binding properties.

Apremilast induced phosphorylation of plakoglobin at serine 665 – a mechanisms which is known to stabilize cardiomyocyte cohesion. Interestingly, keratinocytes phospho-deficient at this side showed altered organization of Dsg1, Dsg3 and keratin filaments and impaired adhesion, which was not rescued by apremilast. These data identified a new mechanism of desmosome regulation and propose that apremilast is protective in pemphigus by stabilizing keratinocyte cohesion.

## Introduction

Epidermal integrity is critically dependent on desmosomes which, as intercellular junctions, are required to maintain proper adhesion and to precisely regulate variable signaling pathways to determine cellular behavior, differentiation and adaption to respective biological requirements ^1,2^. The adhesive function of desmosomes is maintained by the desmosomal cadherins of the desmoglein (Dsg) and desmocollin (Dsc) subfamilies, which are linked via several plaque proteins including plakoglobin (Pg), plakophilins (Pkp) and desmoplakin (Dp) to the intermediate filament cytoskeleton and thus to keratins in the epidermis ^3,4^.

Pemphigus is a life-threatening bullous autoimmune disease in which autoantibodies primarily directed against Dsg1 and Dsg3 cause flaccid blistering in the epidermis and in mucous membranes of the oral cavity and elsewhere ^5,6^. Therapeutic strategies range from unspecific immunosuppression and targeted immunotherapy such as immune-apheresis and rituximab to deplete autoantibody-producing B cells to experimental methods which are not yet clinically approved ^6–13^. All these strategies modulate the immune system of the patients and thus are associated with considerable side effects. Moreover, during the acute phase of the disease until depletion of autoantibodies is accomplished, additional approaches directly stabilizing desmosomal adhesion of keratinocytes would fulfill an unmet medical need. Therefore, a precise understanding of the mechanisms leading to loss of intercellular adhesion in keratinocytes is of high clinical impact. Reported mechanisms comprise direct inhibition of Dsg3 interaction and dysregulation of various intracellular signaling pathways upon autoantibody binding, which finally cause alterations of the keratin cytoskeleton as well as depletion of Dsg molecules from cell membranes of keratinocytes ^14–17^. However, the plethora of signaling mechanisms involved is not yet fully elucidated.

It has been shown that Dsg1 and Dsg3 act as signal transducers and are directly involved in the regulation of specific signaling pathways upon autoantibody binding by formation of signaling complexes with p38MAPK, PI4 kinase, PLC and PKC ^18,19^. Signaling pathways can be differentiated into those contributing directly to the loss of intercellular adhesion and others representing cellular rescue mechanisms. In the first group, signaling pathways such as p38MAPK, PLC, Erk, ADAM10 and Src can be highlighted which are activated upon autoantibody binding and contribute to keratin alterations and Dsg depletion. Accordingly, mediators blocking activation of these signaling pathways can prevent pemphigus-IgG-induced loss of intercellular adhesion *in vitro* as well as *in* and *ex vivo* ^20–25^. In the second group, we previously reported cAMP as a protective signaling pathway in pemphigus ^26^, which is known to strengthen cadherin-mediated adhesion in different tissues as shown for classical cadherins in the endothelium ^27^ and for desmosomal cadherins in the heart. In the myocardium, cAMP induced a phenomenon referred to as positive adhesiotropy, which strengthens intercellular adhesion via phosphorylation of Pg at S665 ^28,29^. In pemphigus, autoantibodies increase cAMP in keratinocytes, which represents a sufficient rescue mechanism to prevent loss of intercellular adhesion when pharmacologically augmented by drugs such as the adenylate cyclase activator forskolin combined with the phosphodiesterase inhibitor rolipram (F/R) or by the β-adrenergic mediators such as isoprenaline ^26^. However, these drugs stimulate adrenergic signaling not only in keratinocytes and thus are not suitable for application in patients. During the past years several more specific phosphodiesterase 4 inhibitors were developed and clinically applied ^30,31^. Among them, apremilast is clinically approved for treatment of psoriasis and Behcet disease and for psoriasis effects on epidermal clinical presentation were reported ^32–35^. Recently, a pemphigus patient has been successfully treated with apremilast ^36^. However, therapeutic effects of apremilast were mainly attributed to the immune system and effects on keratinocytes have not yet been reported. Therefore, we here analyzed the effect of apremilast on desmosomal adhesion and its therapeutic role in pemphigus.

Our data demonstrate that apremilast is effective to prevent PV-IgG-induced blistering in human skin by stabilizing keratinocyte adhesion. This protective effect was paralleled by inhibition of keratin filament retraction for which Pg-phosphorylation at S665 is required. Thus, apremilast may serve as a treatment option during the acute phase in pemphigus.

## Results

### Apremilast is protective in an *ex vivo* pemphigus skin model

Increase of keratinocyte cAMP levels by F/R was shown to protect keratinocytes from PV-IgG-induced loss of intercellular adhesion *in vitro* and isoprenaline also was protective against blister formation in an *in vivo* pemphigus neonatal mouse model ^26^. However, these mediators are inappropriate for application in patients due to their expected systemic side effects. Thus, we here used the phosphodiesterase inhibitor apremilast, which is already clinically approved for the treatment of psoriasis and Behcet disease, and analyzed its effect on impaired desmosomal adhesion in PV. First, we performed a dose kinetic in a dispase-based keratinocyte dissociation assay and compared apremilast powder and tablets, both of which increased intercellular adhesion in HaCaTs in a dose-dependent manner (data not shown). The concentration of 100 μM showed the best effect on cell cohesion and significantly elevated cAMP levels upon treatment for 2 h and 24 h and was used for all subsequent experiments (Figure S1A, B, Figure 2A).

Next, we applied apremilast in a human *ex vivo* skin model. Healthy human skin from body donors was injected subepidermally with apremilast (Apr) or its vehicle (DMSO) followed by either control IgG (C-IgG) or PV1-IgG injection 1 h later. Samples were incubated for 24 h and subsequently subjected to mechanical stress. HE staining of C-IgG-injected samples with and without apremilast showed no blister formation or morphological alteration in the epidermis (Figure 1A, B). In contrast, PV1-IgG injection induced suprabasal blistering with a mean cleft length of 860 μm (+/-220 μm), which was completely abrogated by injection of apremilast indicating that apremilast is effective to prevent blister formation independent of effects on immune cells and thus, most likely by ameliorating loss of intercellular adhesion in keratinocytes (Figure 1A, B).

**Figure 1:**
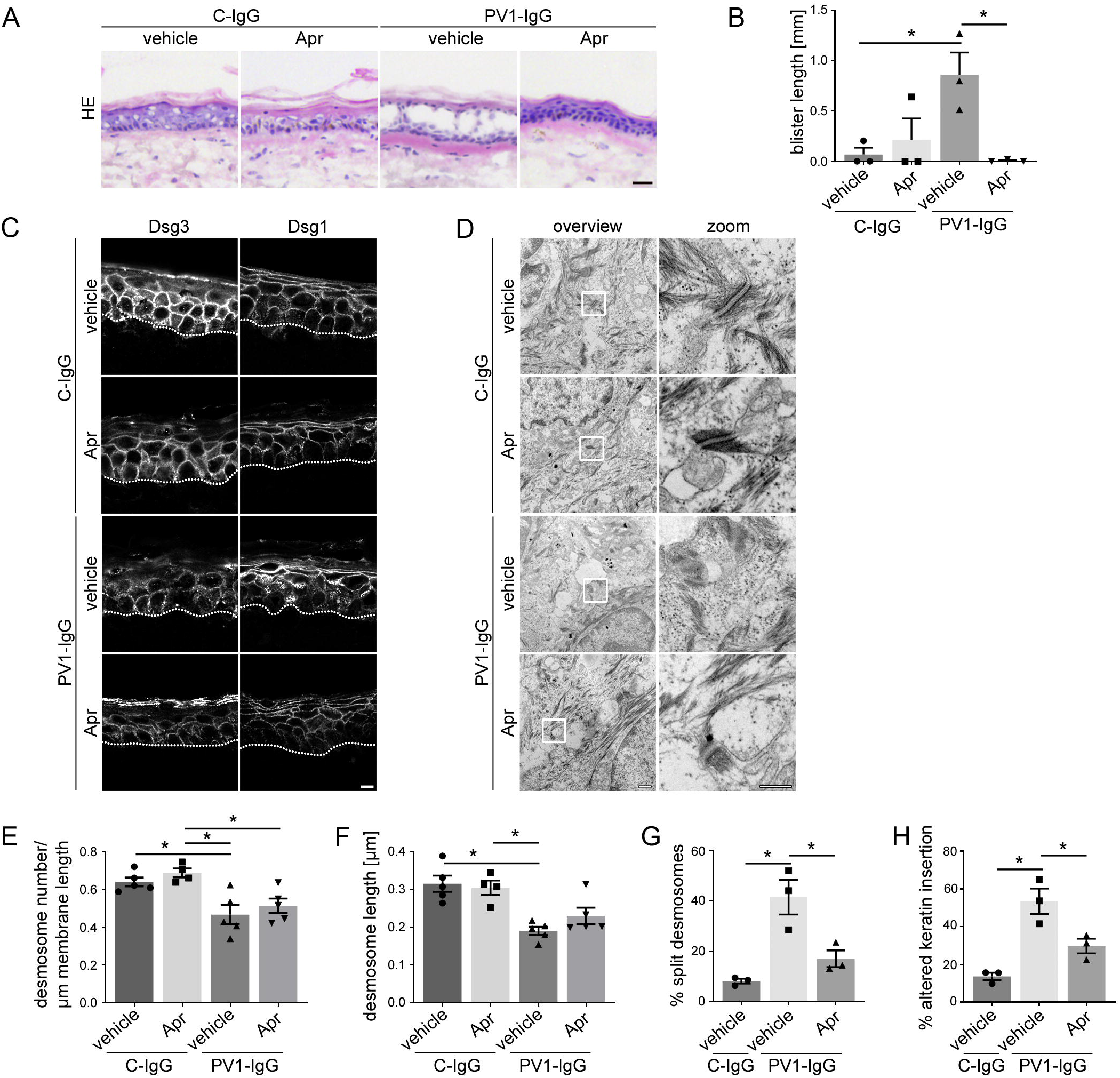
Apremilast is protective in an *ex vivo* pemphigus skin model. (A) HE staining of human *ex vivo* skin samples after injection of vehicle (DMSO) or apremilast (Apr) prior to injection of PVl-IgG or control(C)-IgG from healthy humans. Apremilast protects from PVl-IgG-induced intraepidermal blister formation. Scale bar = 25 μm. Representative of n=3. (B) Quantification of blister length revealed a significant decrease in blister length after apremilast treatment (n=3). (C) Immunostaining against Dsg1, Dsg3 in *ex vivo* skin. Apremilast did not improve PVl-IgG-induced Dsg depletion. Scale bar = 7.5 μm. Representative of n=4. (D) Electron microscopic analysis of desmosome ultrastructure of ex vivo skin samples. Scale bar (overview) = 500 nm. Zoom in areas marked with white rectangles. Scale bar (zoom) = 200 nm. Representative of n=5. Quantification revealed no rescue of PV1-IgG-induced reduction in number (E) and length (F) of desmosomes but showed improvement of PV1-IgG-induced formation of split desmosomes (G) and altered keratin network insertion (H) by apremilast (n=3-5). Columns indicate mean value ÷ SEM, *P<0.05; One-way ANOVA with Bonferroni correction. Desmoglein (Dsg), pemphigus vulgaris (P)

Next, we stained the *ex vivo* samples for PV antigens Dsg1 and Dsg3. The epidermis showed normal inverse expression pattern of Dsg1 and Dsg3 in C-IgG-treated samples (Figure 1C). In contrast, PV1-IgG injection led to a fragmented and reduced Dsg1 and Dsg3 staining in the epidermis, further referred to as Dsg depletion as described before ^37^. Surprisingly, apremilast did not attenuate PV1-IgG-induced Dsg depletion. Specificity of staining was shown by secondary antibody control (Figure S1C).

We next analyzed desmosome ultrastructure of *ex vivo* samples using electron microscopy to investigate the protective mechanisms underlying the apremilast effect. Studies on PV patients’ skin biopsies or PV mouse and human *ex-vivo* models revealed an ultrastructural PV-phenotype characterized by reduction of desmosome number and size, uncoupling of keratins from the desmosomal plaque and desmosomes with separated plaques, referred to as split desmosomes ^38–42^. PV1-IgG-injected *ex vivo* samples showed a reduction in desmosome number and size compared to control samples, which was not rescued by apremilast injection (Figure 1D-F). This is in line with the immunostaining data of Dsg1 and Dsg3 (Figure 1C). In contrast, PV1-IgG-induced formation of split desmosomes was reduced by apremilast treatment (Figure 1D, G), suggesting that apremilast strengthens interaction of the remaining desmosomes. Keratin uncoupling is a morphological hallmark in pemphigus contributing to destabilization of desmosomes in PV. Vice versa, blocking of keratin retraction was shown to be protective against PV-IgG-induced loss of intercellular adhesion ^20,42–45^. Thus, we analyzed the insertion of keratin filaments to the desmosome, which was altered upon autoantibody injection (Figure 1D, H). Importantly, apremilast significantly ameliorated PV-IgG-induced uncoupling of keratins from the desmosome (Figure 1D, H). Taken together, these data suggest that the protective effect of apremilast in PV may be based on improved keratin insertion rather than diminished Dsg depletion.

### Apremilast ameliorates PV-IgG induced loss of cell adhesion and keratin alterations *in vitro*

Next, we analyzed intercellular adhesion of in human keratinocytes (HaCaT) *in vitro*. AK23, a pathogenic anti-Dsg3 antibody derived from a pemphigus mouse model ^46^ as well as PV1-IgG and PV2-IgG impaired intercellular adhesion in HaCaT cells (Figure 2A-C). Pre-treatment with apremilast significantly diminished autoantibody-induced loss of cell cohesion (Figure 2A-C). Further, we repeated these experiments in primary human keratinocytes (NHEK) (Figure 2D). In concordance to results in HaCaT cells, apremilast was sufficient to ameliorate PV2-IgG-induced loss of intercellular adhesion in NHEKs.

**Figure 2:**
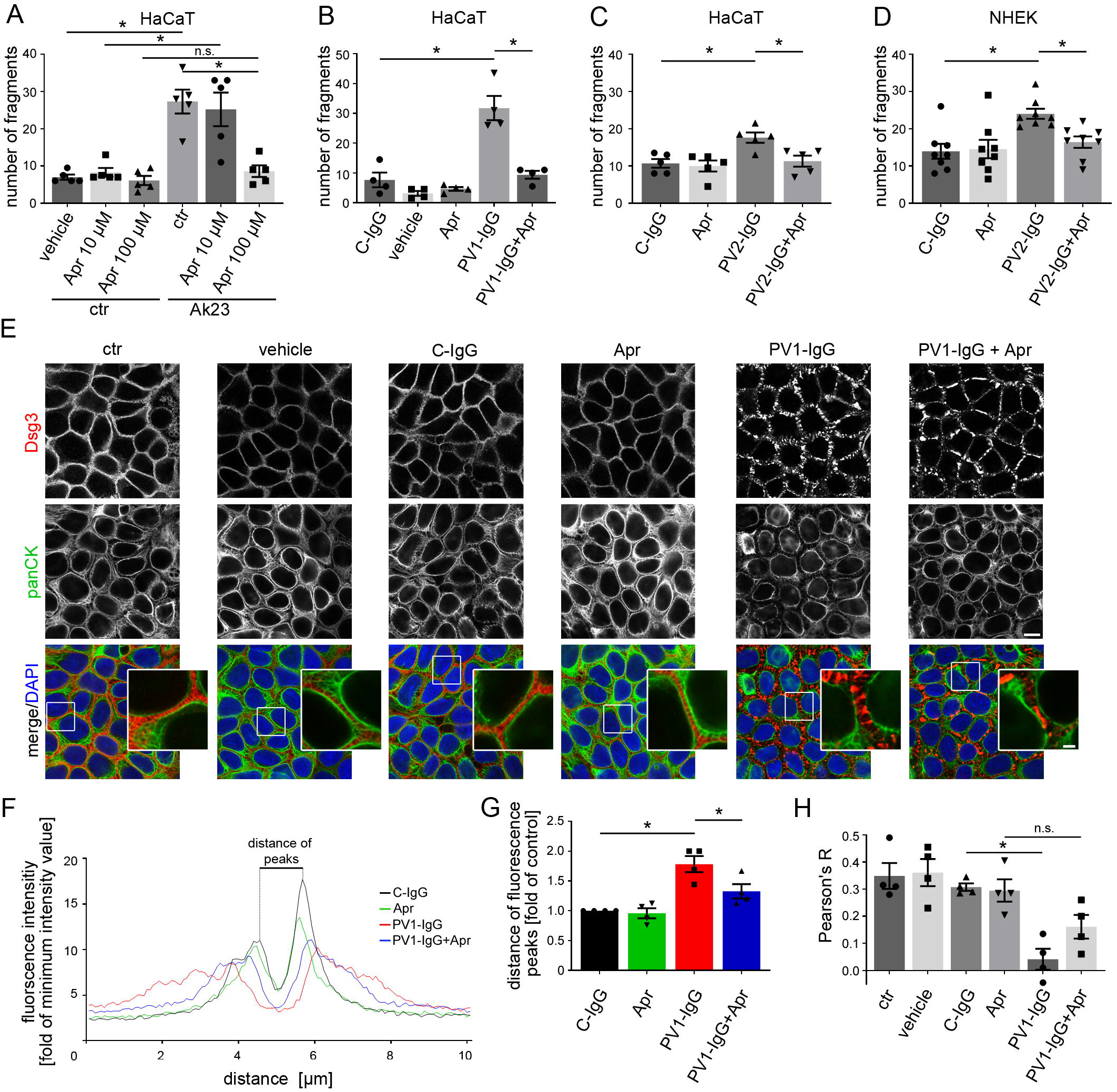
Apremilast ameliorates PV-IgG-induced loss of intercellular adhesion and keratin filament alterations. (A-D) Dispase-based keratinocyte dissociation assay of human keratinocytes. 1 h pre-incubation of apremilast attenuates anti-Dsg3 antibody (AK23) in a dose-dependent fashion (A) as well as PV1-(B) or PV2-IgG-(C) induced loss of cell cohesion in HaCaT cells. (D) Primary keratinocytes (NHEK) show same results with PV2-IgG. Columns indicate mean value ÷ SEM, *P<0.05; One-way ANOVA with Bonferroni correction (n=4-8). (E) Immunostaining of Dsg3 and keratin filaments (panCK) in HaCaTs. Apremilast ameliorates PV 1 -induced keratin retraction but not Dsg3 depletion and fragmentation of staining. Scale bar = 10 μm. Zoom in areas marked with white rectangles. Scale bar (zoom) = 2.5 μm. Representative of n=4. (F, G) Quantification of cytokeratin fluorescence intensity in small areas perpendicular to the respective cell border. (F) Average of keratin fluorescence intensity measured along 10 μm spanning a cell border under respective conditions. (G) Apremilast improved PVl-IgG-induced increase in distance of fluorescence peaks as a measure for retraction of the keratin cytoskeleton (n=4). (H) Calculation of Pearson’s correlation coefficient. Apremilast ameliorates PVl-induced loss of co-localization between keratin filaments and Dsg3 (n=4).

To investigate the effects of apremilast upon PV-IgG treatment on Dsg and keratin organization in more detail, we performed immunostaining against Dsg3 and keratin filaments in HaCaT keratinocytes (Figure 2E). Under control conditions, Dsg3 revealed a linear staining along cell borders and a dense keratin network covered the cytoplasm. PV1-IgG induced fragmentation of Dsg3 staining. Moreover, cytokeratin staining was less intense and retracted from cell periphery similar as described before ^43,44,47^. In line with *ex vivo* data, apremilast did not rescue PV1-IgG-induced Dsg3 depletion (Figure 2E). Same results were yielded with PV2-IgG and also for Dsg1 and 3 in NHEK cells (Figure S2A, B). However, apremilast alone induced condensation of keratin network and thickening of bundles and also ameliorated PV1-IgG-induced keratin retraction (Figure 2E-G). Quantification of cytokeratin fluorescence intensity in arrays perpendicular to the cell border revealed decreased fluorescence intensity and a significantly increased distance between fluorescence peaks in PV-IgG-treated samples which was ameliorated by apremilast (Figure 2F, G). In addition, we calculated Pearson’s correlation coefficient to measure co-localization between keratin filaments and Dsg3. PV1-IgG diminished co-localization between these two proteins, which was rescued by pretreatment with apremilast (Figure 2E, H). Western blot analysis of HaCaT cell lysates supported the finding that apremilast had no effect on PV1-IgG-induced Dsg3 depletion which is known to primarily occur in the triton soluble fraction ^48^ (Figure S2C, D).

Similar results were obtained by analyses of HaCaT cells stably expressing CK5-YFP (Figure 3A). Intercellular contacts in HaCaTs are located in finger-like cell processes containing keratin filaments ^49^. We analyzed the keratin-bearing processes over a 10 μm length of the membrane as a measure of desmosomes anchored to the keratin cytoskeleton. Number of keratin-bearing processes was significantly reduced in PV1-IgG samples (Figure 3A, B). Apremilast treatment completely restored number of keratin-bearing cell processes (Figure 3A, B). Specificity of staining was shown by secondary antibody control (Figure S2E). Previous studies from our group showed Dsg3 single molecule binding properties to be dependent on keratins and that increased cAMP in cardiomyocytes causes stronger adhesion of Dsg2 along cell junctions ^28,50^. Therefore, we wondered if strengthened keratin network and improved Dsg3-keratin colocalization by apremilast would affect Dsg3 binding properties in keratinocytes and performed atomic force microscopy (AFM) adhesion measurements on living HaCaT cells using Dsg3-functionalized tips. Keratin filaments were readily detectable as finger-like processes along intercellular contacts in AFM topography images (Figure 3C, topography overview) similar as described before ^51^. Nevertheless, apremilast did not change Dsg3 binding frequency or binding strength, referred to as unbinding force (Figure 3C-E). Furthermore, step position, which was suggested to be a measure for cytoskeletal anchorage ^52^, was not changed upon apremilast application when no pathogenic autoantibodies were present (Figure 3F). Experiments using apremilast in addition to PV-IgG were not feasible because pemphigus autoantibodies directly interfere with Dsg3 binding in living keratinocytes as shown before ^49^.

**Figure 3:**
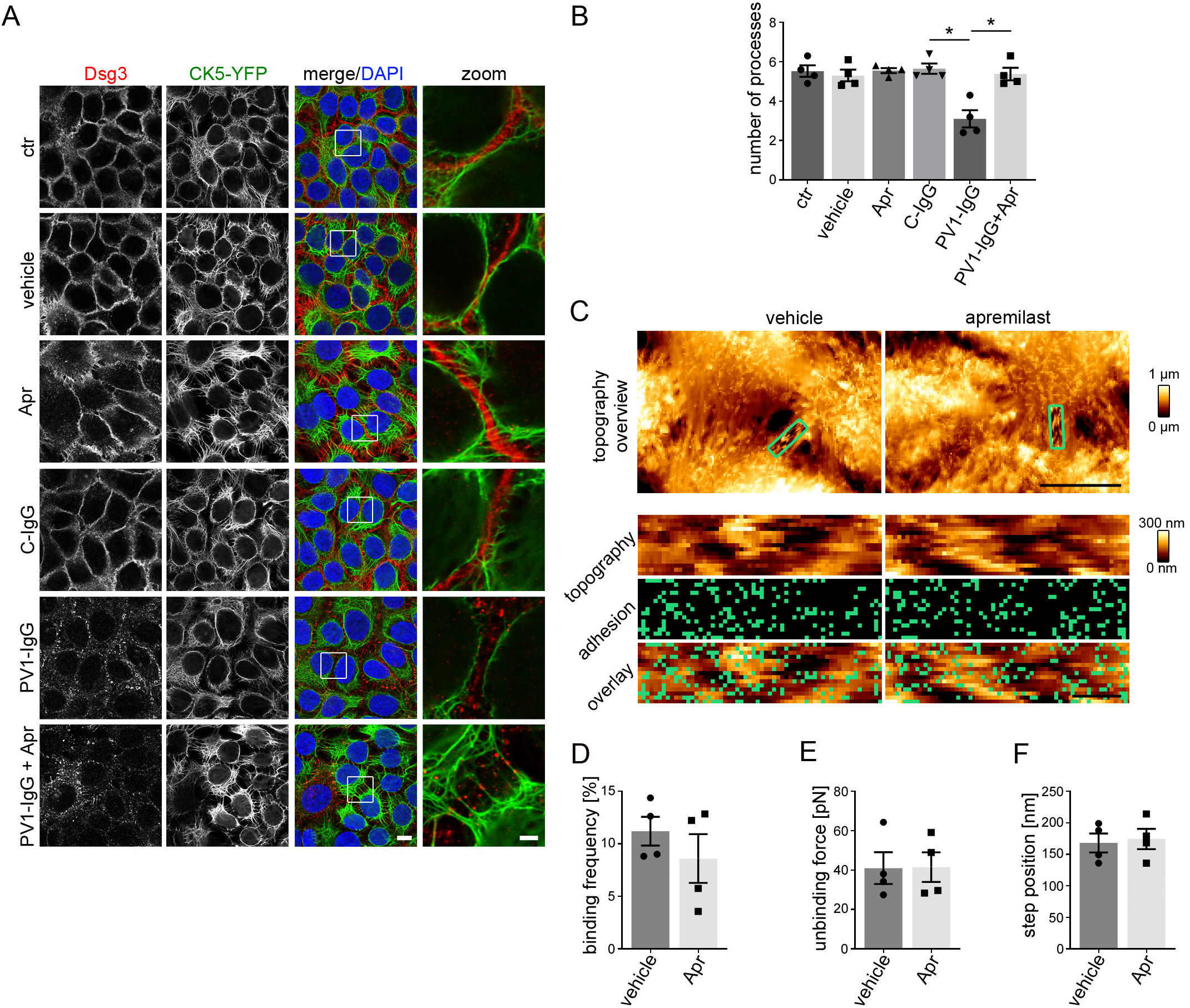
Apremilast restores PVl-IgG-induced retraction of the keratin cytoskeleton but had no effect on Dsg3 alterations. (A) Immunostaining of Dsg3 in stably expressing CK5-YFP HaCaTs. Apremilast prevents PV-IgG-induced retraction of the keratin cytoskeleton but not Dsg3 depletion and fragmentation of staining. Representative of n=4. Scale bar = 10 μm. Areas for zoom in marked with white rectangles. Scale bar (zoom) = 2.5 μm. (B) Quantification of the number of keratin-bearing processes over 10 μm of the membrane. Pretreatment of apremilast restores PVl-IgG-induced decreased number of keratin-bearing cell processes (n=4). (C) Topography overview images of atomic force microscopy (AFM) measurements on living HaCaTs using a Dsg3-functionalized tip revealing cell borders bridged by dense filamental structures. Scale bar = 10 μm. Small areas along the cell borders (green rectangles) were chosen for adhesion measurements. In these areas each pixel represents a force-distance-curve. In the adhesion panel each green pixel represents a Dsg3-specific binding event. Scale bar = 1 μm. (D-F) Quantification of AFM adhesion measurements. (D) Apremilast had no effect on Dsg3 binding frequency. Additionally, the unbinding force (E) as a measure for the single molecule binding strength as well as the step position (F) as a measure for cytoskeletal anchorage of the measured molecules were unaltered in cells treated with apremilast (8 cell borders from 4 independent experiments with 900 force-distance curves/ cell border). Columns indicate mean value ± SEM. *P<0.05.

In accordance with the *ex vivo* data, these results reveal that apremilast strengthens cell contacts by stabilization of the keratin network rather than by affecting Dsg3 turnover and adhesive properties.

### Apremilast-induced phosphorylation of plakoglobin at S665 is required for keratin filament organization in keratinocytes

In cardiomyocytes cAMP-dependent phosphorylation of Pg at S665, a mechanism, referred to as positive adhesiotropy, strengthens cell cohesion ^28^. Using a phospho-specific antibody for Pg-S665 ^29^, we observed that in intact human epidermis apremilast increases localization of pPg (S665) along keratinocyte cell membranes (Figure 4A). Furthermore, apremilast induced significant Pg phosphorylation at S665 in HaCaTs compared to control conditions as revealed by Western blot analysis (Figure 4B, C).

**Figure 4:**
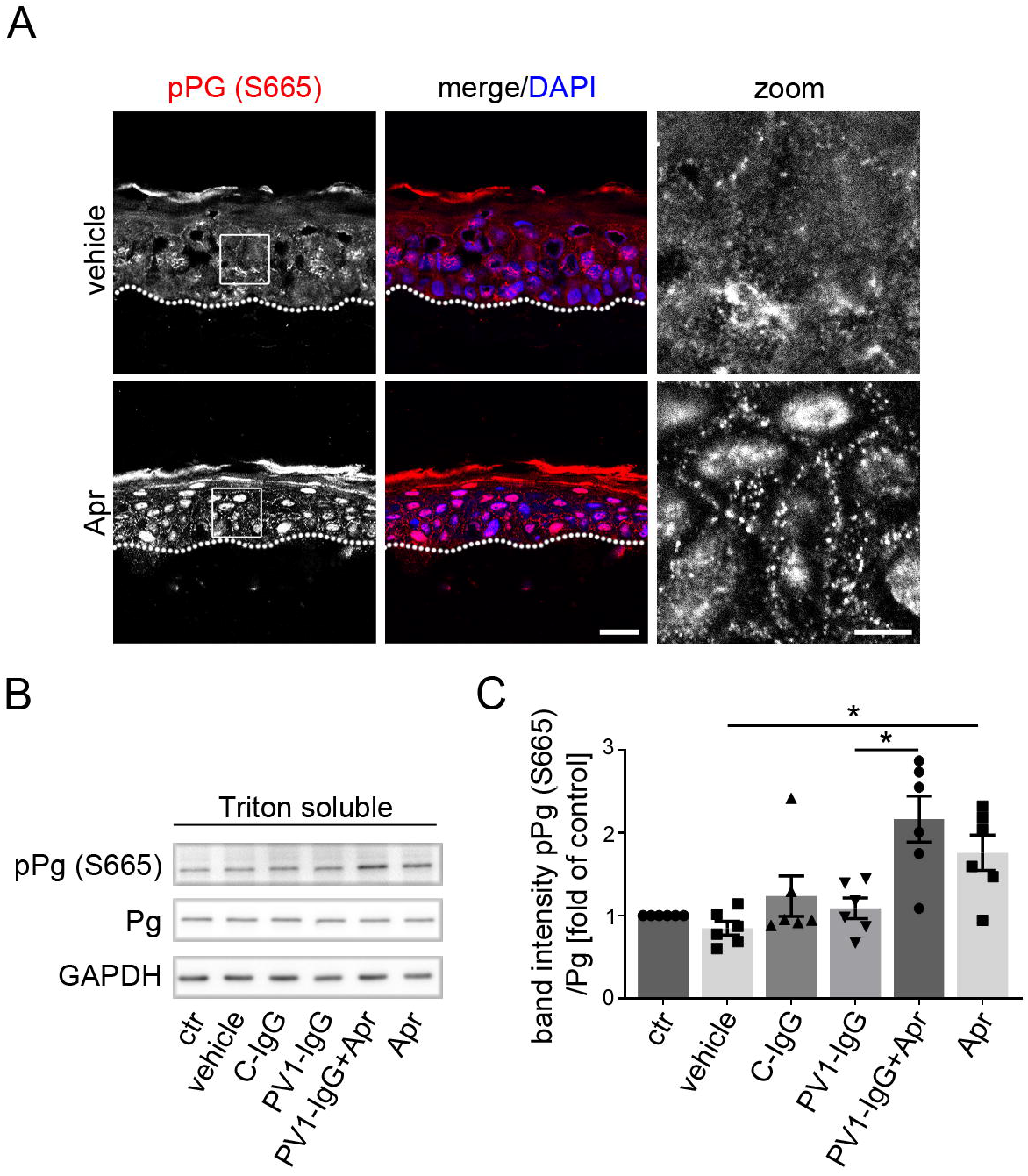
Apremilast increases phosphorylation of Pg at S665. (A) Immunostaining of phosphorylated Pg at S665 in human ex vivo skin samples shows an increased phosphorylation of Pg at S665 upon apremilast injection (n=4). Scale bar = 20 μm, zoom in marked by a white rectangle. Scale bar (zoom in) = 5 μm. Representative Western blot of HaCaT Triton soluble fraction (B) and quantification ofband intensity ofpPg (S5665) (C) show a significant increased phosphorylation of Pg at S665 after apremilast treatment (n=6). GAPDH and total Pg was used as loading control. Columns indicate mean value ± SEM. *P<0.05. One-way ANOVA with Bonferroni correction.

### Phosphorylation of plakoglobin at S665 is required for proper desmosomal adhesion and keratin organization in keratinocytes of murine epidermis

The data above suggest a similar mechanism to positive adhesiotropy in cardiomyocytes via cAMP-mediated Pg phosphorylation at S665 that contributes to strengthening of intercellular adhesion to be present in keratinocytes. Thus, we generated a phospho-deficient Pg-S665A *knock in* mouse model, where serine 665 was replaced by alanine (Figure S4D). Homozygous Pg-S665A mice were viable and did not show significant macroscopic alterations in internal organs (data not shown). For further investigation on the effect of Pg phosphorylation at S665 for keratinocyte adhesion we generated two murine keratinocyte cell lines from both wt (wt-1/-2) and Pg phospho-deficient mice (Pg-S665A-1/-2), respectively. First, immunostainings of desmosomal proteins were performed. Interestingly, cells bearing the phospho-deficient mutant of Pg (Pg-S665A) were smaller in size and only barely differentiated in 2-3 cell layers as typical for wt murine keratinocytes (Figure 5A). In accordance, differentiation-dependent expression of Dsg1 was drastically altered. Color-encoded 3D-representation of wt keratinocytes revealed that Dsg1 was mainly present in the superficial cell layers as expected from Dsg1 distribution in intact epidermis. In contrast, Dsg1 was reduced and in large parts of the monolayer only one cell layer was present in Pg-S665A keratinocytes (Figure 5A, B) demonstrating the importance of Pg phosphorylation for both turnover of desmosomal proteins and differentiation of the keratinocytes. Comparable to the Z-stacks Dsg1 appeared fragmented and with a reduced intensity in Pg-S665A keratinocytes (Figure 5B). Interestingly, in wt keratinocytes desmoplakin (Dp) was distributed linearly along cell borders. In contrast, Dp revealed a dotted localization pattern in Pg-S665A keratinocytes arguing for alterations in desmosomal turnover in these cells (Figure 5B). Similarly, Dsg3 staining was reduced and fragmented in Pg-S665A keratinocytes (Figure S3A). Further, Pg was slightly reduced along cell borders and pPg (S665) was absent along cell borders in phospho-deficient keratinocytes (Figure 5C, D).

**Figure 5:**
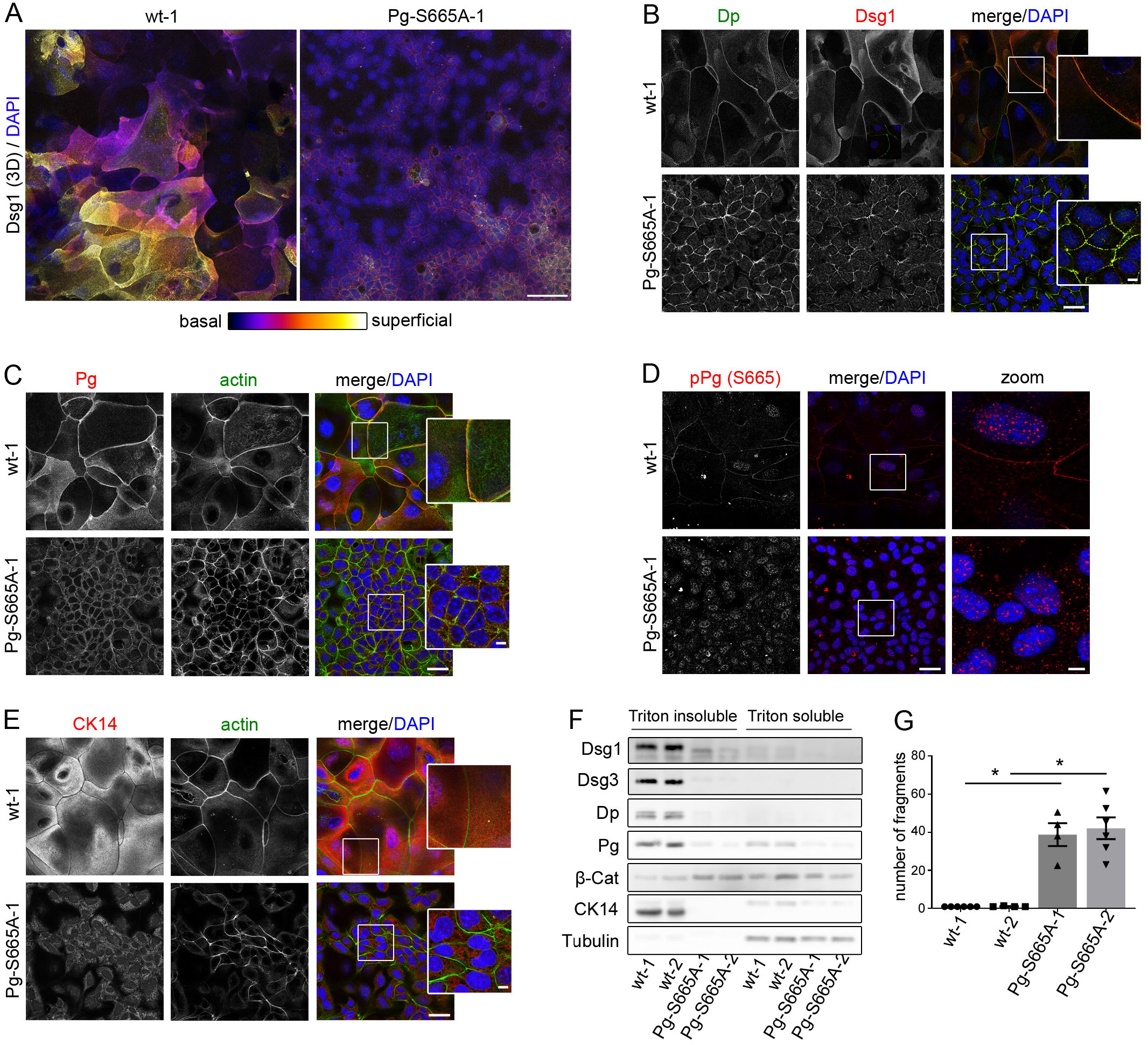
Keratinocytes phospho-deficient at Pg S665 (Pg-S665A) reveal impaired intercellular adhesion accompanied with drastic changes in desmosomal proteins and keratin filament cytoskeleton. (A) Color-encoded Z-stack of wt and Pg-S665A keratinocytes reveal multiple layers in wt and only 1-2 layers in Pg-S665A keratinocytes. Representative of n=4. Scale bar = 50 μm. (A, B) Dsgl is mainly expressed in the superficial layers in a linear fashion along cell borders. In contrast, Dsgl is drastically reduced and restricted to the basal layer in Pg-S665A keratinocytes. (B) Dp is slightly reduced and more dotted along the cell borders in Pg-S665A keratinocytes in comparison to wt. (C) Interestingly, Pg is reduces and appears less linearized in Pg-S665A keratinocytes compared to wt whereas the actin cytoskeleton is broadened in Pg-S665A. (D) In accordance with murine epidermis, pPg-S665 is absent in Pg-S665A keratinocytes confirming their phospho deficiency at this site whereas it appears dotted along cell borders and in the nuclei in wt keratinocytes. (E) The keratin cytoskeleton shows a dense network spanning the whole cell in wt keratinocytes whereas it is rarefied and drastically altered in Pg-S665A keratinocytes. (B-E) Representatives of n=3-4. Scale bar = 25 μm. White rectangles depict areas for zoom in. Scale bar (zoom) = 5 μm. (F) Triton-based separation reveals a drastic impairment of Dsgl and 3 expression in the Triton insoluble, desmosome bearing fraction in Pg-S665A keratinocytes. Similar observations were made for Dp and Pg. Importantly, keratin 14 (CK14) is almost absent in Pg-S665A keratinocytes. In contrast, ß-catenin was slightly upregulated in the Triton insoluble fraction. Tubulin serves as loading control. Representative of n=4. (G) Dissociation assay in two different wt and Pg-S665A keratinocyte cell lines reveal a drastic impairment of intercellular adhesion in cells with phospho-deficient Pg at S665. n= 4-5. Columns indicate mean value ÷ SEM. *P<0.05. One-way ANOVA with Bonferroni correction.

Importantly, the keratin network which builds up a dense and delicate mesh throughout the whole cytoplasm in wt keratinocytes was substantially altered in Pg-S665A keratinocytes and appeared thinned out and irregular without a clear mesh formation (Figure 5E). These data demonstrate that Pg phosphorylation at S665 is critically involved in proper keratin filament organization. We further observed actin bundles to be broadened and more prominent, which may serve as a compensatory mechanism in Pg-S665A keratinocytes (Figure 5C, E). Along the same line, the adherens junction proteins β-catenin and E-cadherin were upregulated (Figure S3B, C). Interestingly, mRNA levels of *Pg* were unaltered, indicating that impaired Pg localization along cell borders is independent mRNA levels (Figure S3D). To confirm these results, we performed Western blot analysis of the respective proteins in two wt and Pg-S665A cell lines. Indeed, Dsg1 and 3 as well as Pg and Dp were reduced especially in the triton-insoluble and thus desmosome-containing fraction (Figure 5F). Further, an almost complete loss of CK14 clearly reflected the importance of Pg phosphorylation at S665 for keratin organization (Figure 5F).

Next, we investigated the effect of Pg-S665A on intercellular adhesion in the respective keratinocytes. Importantly, Pg-S665A keratinocytes showed a significantly and drastically impaired intercellular adhesion as revealed by dissociation assays (Figure 5G).

Subsequently, we asked whether a pathogenic pemphigus autoantibody such as AK23 would have further effects on cell adhesion in Pg-S665A keratinocytes and, if so, protective effects of apremilast were present. We applied AK23 and apremilast for 24 h in wt and Pg-S665A keratinocytes and performed dissociation assays. In wt cells, AK23 led to fragmentation and thus loss of intercellular adhesion which was significantly reduced by apremilast (Figure 6A). In contrast, the additional effect of AK23 on weakened intercellular adhesion in Pg-S665A was not significant. Further, no protective effect of apremilast was observed in the absence or presence of AK23, indicating that protective effects of apremilast in pemphigus are dependent on Pg phosphorylation at S665 (Figure 6A).

**Figure 6:**
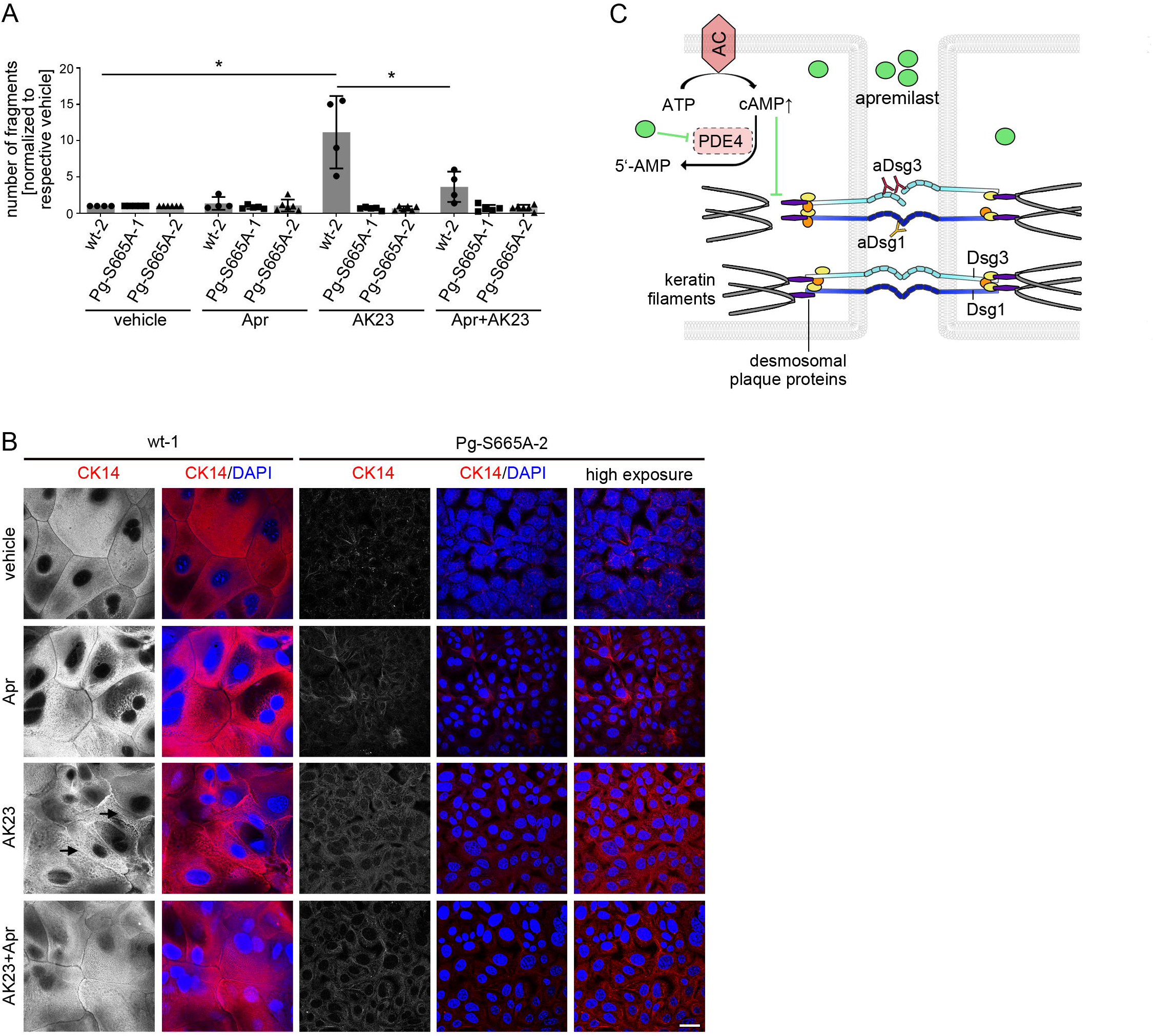
In Pg-S665A keratinocytes, a pemphigus autoantibody and apremilast had no effects on intercellular adhesion. (A) Dissociation assay in wt murine keratinocytes reveal a significant fragmentation after 24 h of incubation with the pathogenic monoclonal Dsg3 antibody AK23. Pre-incubation with apremilast (Apr) for 1 h ameliorates AK23-induced loss of intercellular adhesion in wt keratinocytes. In contrast, in Pg-S665A keratinocytes, which already show a drastic impairment of intercellular adhesion in untreated conditions, AK23 had no additional effect on intercellular adhesion. Apremilast did not restore impaired intercellular adhesion neither under basal conditions nor after AK23 incubation, n=3-5. Columns indicate mean value ± SEM. P<0.05. One-way ANOVA with Bonferroni correction. (B) Keratin 14 (CK14) staining in wt and Pg-S665A murine keratinocytes under same conditions as dissociation assay. In wt, keratin cytoskeleton forms a dense network spanning the whole cytoplasm under control condition. AK23 leads to keratin alterations with coarsened keratin bundles (arrows) which were completely abolished by pre-treatment with apremilast. In Pg-S665A keratinocytes the keratin cytoskeleton is rarefied and drastically altered under basal conditions. AK23 led to a mild increased irregularity although effects were minor due to the drastic phenotype under basal conditions. Apremilast had no significant effect on composition and filament bundle distribution under basal conditions and after treatment with AK23. A high exposure panel was included for better visibility of the keratin cytoskeleton in Pg-S665A keratinocytes. Representative of n=3. Scale bar = 25 μm. (C) Apremilast prevents PV-IgG induced loss of intercellular adhesion. Autoantibodies directed against desmogleins induce uncoupling of keratin filaments from the desmosomal plaque. Increase of intercellular cAMP by the phosphodiesterase inhibitor apremilast ameliorates this effect and thereby rescues loss of cell adhesion.

Apremilast strongly inhibited PV-IgG-induced keratin alterations *ex vivo* and in human keratinocytes. Thus, we next performed immunostaining for CK14 in wt and Pg-S665A keratinocytes under the same conditions. In wt cells, AK23 led to a coarsening of the keratin cytoskeleton and reduction of keratins especially in the cell periphery which was in part blocked by apremilast treatment (Figure 6B, arrows). In contrast, as expected, no additional alterations of keratin organization were observed in Pg-S665A keratinocytes when treated with apremilast or AK23 alone or in combination.

Taken together, these data demonstrate Pg phosphorylation at S665 to be crucial for proper adhesive function of desmosomes via its contribution to proper organization of the keratin cytoskeleton of keratinocytes.

## Discussion

Pemphigus is a group of severe autoimmune disease treated by immunosuppressive drugs and especially rituximab to deplete autoantibody-producing B cells. However, the time gap until the immunosuppressive medication is sufficient to ameliorate patients’ symptoms can at present only be bridged by corticosteroids ^5,6,13^. These therapeutic strategies do all directly target the immune system and as such, are associated with severe side-effects. In contrast, no medication is available that directly targets adhesion of human keratinocytes. Therefore, we characterized mechanisms of protective cAMP signaling to strengthen desmosomal adhesion in keratinocytes upon binding of anti-Dsg IgG and investigated apremilast as a new therapeutic strategy in PV, the main pemphigus variant. Indeed, apremilast increased cAMP levels in keratinocytes and effectively prevented both PV-IgG-induced loss of desmosomal adhesion *in vitro* and acantholysis in human epidermis *ex vivo*. Importantly, apremilast blocked keratin filament reorganization, which is established to be a morphological hallmark in pemphigus present in both patients’ skin lesions and human skin models ^42,53^. Using a new Pg-S665-phosphodeficient mouse model we found that phosphorylation of Pg at S665 is crucial for maintenance of keratin organization and keratinocyte adhesion.

### Apremilast induces protective cAMP signaling in pemphigus

Previously, cAMP increase was shown to regulate cadherin-mediated adhesion in various tissues. For instance, for VE-cadherin, cAMP increase was reported both to stabilize VE-cadherin at the cell membrane and to contribute to proper barrier function of the endothelium ^54^. Further, cAMP increase in cardiomyocytes led to a phenomenon referred to as positive adhesiotropy and had a direct effect on adhesive strength of cardiac desmosomes. Adrenergic signaling induced redistribution of the desmosomal cadherin Dsg2 towards cell borders of cardiomyocytes and strengthened intercalated discs on the ultrastructural level ^28,29^.

Here, we show that apremilast via protective cAMP signaling in keratinocytes is a promising approach to target loss of desmosomal adhesion in pemphigus patients. Previously, we reported that cAMP increase induced by the combination of the adenylatcyclase (AC) activator forskolin with the phosphodiesterase inhibitor rolipram (F/R) similar to the β-sympathomimetic isoprenaline abolished loss of desmosomal adhesion caused by pemphigus autoantibodies *in vitro* and in a passive immune transfer mouse model *in vivo* ^26^.

Several signaling mechanisms were shown to be critically involved in pemphigus pathogenesis, including p38MAPK, ERK, Src, PKC and Ca^2+^ ^14–16,18^. These signaling pathways are dysregulated upon autoantibody binding and drive loss of intercellular adhesion. Accordingly, modulation of signaling was frequently shown to be protective in different models of pemphigus. However, a therapeutic strategy can only rarely be derived from these investigations as mediators targeting the respective signaling pathways systemically in patients most likely would cause severe side effects. Nevertheless, a recent study identified novel treatment targets including MEK1, TrkA, PI3Kα, and VEGFR2 in pemphigus using an unbiased library approach, revealing the importance of therapies in pemphigus directly targeting keratinocytes ^23^. Therefore, since apremilast is clinically approved to be effective and safe in psoriasis patients ^33^, we further characterized the mechanisms underlying the protective effects of apremilast on keratinocyte adhesion.

Strikingly, apremilast abolished keratin filament reorganization, which is known to be an ultrastructural hallmark in both pemphigus patients’ skin lesions and human skin models (Wilgram 1961; Egu BJD 2017, summarized in Figure 6C). In our experiments, apremilast abrogated retraction of endogenous keratin filaments in cultured cells and intact epidermis *ex vivo* similar to filaments in cells overexpressing CK5-YFP. This observation was also confirmed on the ultrastructural level using transmission electron microscopy in the *ex-vivo* human pemphigus model. Here, changes in keratin insertion in desmosomes after PV-IgG treatment were significantly attenuated by apremilast. These data support previous findings that protective effects in pemphigus models were paralleled frequently by preventing keratin retraction ^43,55^. Moreover, keratins have been demonstrated to critically contribute to the proper adhesive function of desmosomal cadherins and thus to the integrity of desmosomes ^45,50,56^. Interestingly, in contrast to F/R and isoprenaline ^26^, treatment with apremilast did not affect Dsg depletion both *in vitro* and in human skin *ex vivo*. This discrepancy may be explained by distinct amounts of cAMP in keratinocytes treated with either apremilast or F/R but may also be due to different downstream targets or compartmentalization of cAMP ^57^. In line with this observation, keratin retraction and Dsg3 depletion were shown to independently contribute to loss of intercellular adhesion in pemphigus ^44^.

### Plakoglobin S665 phosphorylation is required for keratin organization and intact cell adhesion

In cardiomyocytes positive adhesiotropy was dependent on the desmosomal plaque protein Pg and its phosphorylation at S665 via protein kinase A (PKA). In accordance, overexpression of a phospho-deficient Pg-mutant (Pg-S665A) abolished positive effects of cAMP increase on intercellular adhesion of cardiomyocytes ^28^. Similarly, in keratinocytes PKA was shown to contribute to protective cAMP signaling in pemphigus ^26^. Because we observed that Pg is also phosphorylated at S665 in keratinocytes when cAMP levels were increased by incubation with apremilast, we further investigated the role of Pg S665-phosphorylation in protective cAMP signaling in keratinocytes. To this end, we established a new Pg-S665A-phospho-deficient *knock-in* mouse model and derived murine keratinocytes from these mice for further investigations.

Although mice bearing the Pg-S665A mutation were viable and showed no significant macroscopic alterations in internal organs, desmosomal composition of keratinocytes, derived from these mice was critically altered. The pemphigus antigens Dsg1 and Dsg3 were downregulated and the keratin cytoskeleton was severely compromised. Although the role of Pg in keratin filament organization and keratin retraction in pemphigus was previously reported ^58,59^, the importance of Pg phosphorylation at S665 was not investigated before. Here, we observed that phospho deficiency of Pg at S665 drastically impairs intercellular adhesion of keratinocytes.

### Apremilast-induced cAMP increase in keratinocytes as a therapeutic strategy in skin diseases

Apremilast is a well-established therapeutic approach in psoriasis ^32–34^. However, the mechanisms underlying the clinical effects of apremilast were up to now attributed to modulation of the immune system ^32^. For pemphigus, a recent case report in which a chronic, non-responsive PV patient was successfully treated with apremilast underlines the clinical importance of apremilast in treatment of skin diseases ^36^. Here, we report that PDE4-inhibition by apremilast not only affects the immune system, but also leads to a protective cAMP increase and strengthens intercellular adhesion in keratinocytes. Interestingly, it was reported recently that cAMP levels are increased and AC9 was upregulated in psoriatic skin models ^60^ and that β2-adrenergic receptors are downregulated in psoriatic skin lesions ^61^. Therefore, it was postulated that cAMP may contribute to psoriatic hyperproliferation. However, in light of our observation that increased cAMP may serve as a rescue pathway in pemphigus, the findings may also be interpreted similarly, indicating that positive adhesiotropy induced by adrenergic signaling is a general rescue pathway in inflammatory skin disorders. This hypothesis is further strengthened by the fact that apremilast is an effective and safe therapeutic mediator in psoriasis ^62,63^ which may also have direct effects on keratinocytes.

Interestingly, various case reports showed a simultaneous occurrence of PV and psoriasis ^64,65^ and an elevated risk for pemphigus was suggested for patients suffering from psoriasis ^66^, suggesting a similar predisposition for both diseases by dysregulated cAMP signaling in keratinocytes.

Furthermore, apremilast was shown to be beneficial in a clinical study on palmoplantar keratoderma ^67^, which at least in part may also be effects of apremilast on keratin filament organization in keratinocytes Thus, apremilast-mediated cAMP signaling and its effects on keratinocyte function represents an important therapeutic mechanism of apremilast and opens up a wide range of clinical applications.

Taken together, our data provide a novel therapeutic approach in pemphigus by direct protective cAMP signaling in keratinocytes through apremilast. In addition, we demonstrate a new mechanism involved in desmosome regulation via cAMP-induced Pg phosphorylation at S665. Therefore, we propose a new paradigm according to which apremilast may be effective to treat skin diseases including psoriasis and pemphigus not only by modulation of the immune system but rather also via direct effects on keratinocytes.

## Material and Methods

For extended information please refer to Suppl. Materials & Methods

### Cell Culture and Test Reagents

Experiments were performed in HaCaT immortalized human keratinocytes ^68^, murine keratinocytes or normal human keratinocytes (NHEK). For some experiments, HaCaT cells stably transfected with keratin5-YFP (kind gift of Reinhard Windoffer and Nicole Schwarz, Institute of Molecular and Cellular Anatomy, RWTH Aachen University) were used. HaCaTs were grown in Dulbecco’s Modified Eagle Medium supplemented with 10% FCS (Biochrom, Berlin, Germany), 50 U/ml penicillin and 50g/ml streptomycin (both AppliChem, Darmstadt, Germany) and NHEKs were cultured in epidermal keratinocyte medium (CnT 0.7, Cellntec, Bern, Switzerland). NHEKs were grown in low calcium (Ca^2+^) (0.06 mM) until confluency and switched to 1.8 mM Ca^2+^ 24 h prior experiments. Experiments were performed between passage 3 and 6.

Murine keratinocytes from JUP S665A mice strain were generated as described in detail before ^56,69^. In short, epidermis of neonatal mice were removed using Dispase II (Sigma-Aldrich, Munich, Germany) and epidermal cells were isolated by treatment with Accutase (Sigma-Aldrich) for 1 h. Cells were seeded in complete FAD media (0,05 mM CaCl_2_, PAN Biotech, Aidenbach, Germany) on flasks coated with collagen I (rat tail; BD Bioscience, New Jersey, US). Keratinocytes immortalize spontaneously after 10-15 passages. Subsequently cells were grown under previous mentioned conditions and switched to high Ca^2+^ (1.2 mM) 48 h before experiments were started ^69^.

Cells were maintained in a humified atmosphere with 5% CO_2_ at 37°C (HaCaT, NHEK) or 35°C (murine keratinocytes).

### Purification of patients IgG fractions and AFM proteins

Sera samples of healthy volunteers and PV patients were used with informed and written consent under approval of the local ethic committee (University of Lübeck, AZ12-178). IgG fractions were purified as described before and in Suppl. M&M ^19,70^.

AK23, a pathogenic monoclonal anti-Dsg-3-antibody produced in a mouse model, was bought from Medical & Biological Laboratories Co., Ltd. (Nagoya, Japan) and used at a concentration of 75 μg/ml.

Dsg3-Fc construct for AFM measurements comprising the full extracellular domain of Dsg3 were expressed in Chinese hamster ovary cells. Protein containing supernatants were collected and Protein A-Agarose-based purification was performed as described in detail before ^70^. Measurement of cAMP levels

The cAMP enzyme linked immunosorbent assay (EIA, CA-200, Sigma-Aldrich, St. Louis, MO, USA) was used to evaluate the levels of intracellular cAMP levels. Cells were incubated, washed and lysed by 0.1 M HCl. Afterwards the EIA was performed according to the manufacturer’s manual.

### Keratinocyte dissociation assay

Keratinocytes dissociation assay was performed as described in detail before ^71^. In brief, confluent keratinocyte monolayers were removed from well bottom after respective incubation using 200 μl Dispase II (Sigma Aldrich, St. Louis, Missouri). In murine keratinocytes Dispase II solution was supplemented with 1% of collagenase I (Thermo Fisher Scientific, Massachusetts, US). Subsequently, monolayers were subjected to a defined mechanical stress. Resulting fragments are an inverse measure for intercellular adhesion and were counted either manually or using ImageJ (NIH, Bethesda, Maryland).

### *Ex-vivo* pemphigus skin model

Human *ex vivo* pemphigus model was performed with cadavers of the human body donor program without history of skin diseases from the Institute of Anatomy and Cell Biology, Ludwig-Maximilians-University Munich, Germany as described before ^42^. Written informed consent was obtained from body donors for the use of research samples. Skin samples were taken from body donors who had died no longer than 24 hours before. A 30.5 G needle (B.Braun, Melsungen, Germany) was used to inject either 50 μl Apremilast in DMSO or DMSO and after 1 h of pre-incubation another 50 μl of either PV- or C-IgG. Samples were incubated in Dulbecco’s modified Eagle’s medium (DMEM) in a humidified atmosphere of 95% air and 5% CO_2_ at 37°C for 24 h. After incubation a defined shear stress was applied using a rubber head. Subsequently, specimen were either embedded in Tissue-tec (Leica, Wetzlar, Germany) for cryosectioning or in glutaraldehyde for electron microscopy analysis

### Electron microscopy

For electron microscopy analyses small specimens were fixed in glutaraldehyde, post-fixed in 2% osmium tetroxide and embedded in EPON 812 resin (SERVA Electrophoresis GmBH, Heidelberg, Germany) as described before and in Suppl. M&M ^42^. Specimens were sliced at 60nm thickness using ultramicrotome (Reichert-Jung Ultracut E, Optische Werke AG, Vienna, Austria) and sections were contrasted using uranyl acetate and lead citrate. Images were captured with a Libra 120 transmission electron microscopy (Carl Zeiss NTS GmbH, Germany) equipped with a SSCCD camera system (TRS, Olympus, Tokyo, Japan). For quantification of ultrastructural images please refer to Suppl. M&M.

### Cell lysis and Western blotting

For protein analysis cells were lysed in SDS-lysis-buffer. For some experiments a Triton-X-100-based extraction buffer and centrifugation was used to split triton-soluble non-cytoskeletal bound from the cytoskeletal bound non-soluble proteins. Supernatants containing the Triton-soluble fraction were collected and pellets were resuspended in SDS-lysis-buffer. Electrophoresis and western blot were performed as standard protocols as described elsewhere ^72 73^.

The following primary antibodies were used: anti-Dsg3 rabbit polyclonal (Biozol, Eching, Germany), anti-Dsg1 mouse monoclonal and anti-GAPDH mouse monoclonal (both Santa Cruz, Dallas, TX, USA), anti-plakoglobin mouse (Progen, Heidelberg, Germany), anti-desmoplakin rabbit polyclonal (Abclonal, MA, USA), anti-α-Tubulin and anti-CK14 monoclonal (Abcam, Berlin, Germany), β-catenin monoclonal (BD, Eysins, Switzerland) and anti-p-plakoglobin S665 ^29^.

The following HRP-coupled secondary antibodies have been used: peroxidase-coupled-goat-anti-rabbit or –goat-anti-mouse (Dianova, Hamburg, Germany) was detected using the ECL-system.

### Immunostaining

For immunostaining samples were fixed with PFA or ethanol/aceton, permeabilized if necessary with 0,1 % Triton-X-100 and incubated at 4° with the following primary antibodies: anti-desmoglein-3 rabbit polyclonal (Biozol, Eching, Germany), anti-desmoglein-3 mouse monoclonal (Invitrogen, Carlsbad, CA, USA), anti-desmoglein-1 rabbit polyclonal (Abclonal, MA, USA), anti-Desmoglein-1 mouse monoclonal (Progen, Heidelberg, Germany), anti-desmoglein-2 mouse monoclonal (Origene, Rockville, MD, USA), anti-E-cadherin mouse monoclonal (BD, Eysins, Switzerland), anti-β-catenin mouse monoclonal (BD, Eysins, Switzerland), anti-panCK-FITC mouse monoclonal (Sigma Aldrich, St. Louis, MO), anti-panCK mouse monoclonal (Sigma Aldrich, St. Louis, MO), anti-CK14 mouse monoclonal (Abcam, Berlin, Germany), anti-plakoglobin mouse monoclonal (Progen, Heidelberg, Germany), anti-p-plakoglobin S665 ^29^ and anti-vinculin rabbit polyclonal (Sigma, Aldrich, St. Louis, MO). As secondary antibodies alexa488-, Cy2- or Cy3-coupled goat-anti-rabbit or goat-anti-mouse secondary antibodies (all dianova) were used. Phalloidin Alexa 488 (Molecular Probes, Eugene, OR, USA) was used to stain actin. 4’,6-diamidin-2-phenylindol was added to secondary antibody incubation (DAPI, 1 mg/ml). Afterwards probes were mounted with NPG (1 % n-Propylgallat und 60 % Glycerin in PBS). Images were acquired with a Leica SP5 confocal microscope equipped with a 63× oil objective (Leica, Wetzlar, Germany).

### HE staining

HE staining was performed according to standard procedures and mounted in DEPX (Sigma-Aldrich).

### AFM measurements

AFM measurements on living HaCaT keratinocytes were performed as described before in detail ^49,51^. Briefly, a NanoWizard^®^ 3 AFM (JPK-Instruments, Berlin, Germany) mounted on an inverted optical microscope (Carl Zeiss, Jena, Germany) or Nanowizard 4 AFM (JPK-Instruments, Berlin, Germany) mounted on an inverted optical microscope (IX73 Olympus, Hamburg, Germany), which both allow selection of the scanning area by visualizing the cells with a 63x objective were used. Pyramidal-shaped D-Tips of Si3N4 MLCT cantilevers (Bruker, Mannheim, Germany) were functionalized with Dsg3-Fc as described before ^74^.

AFM was used performed either in Quantitative Imaging (QI, for overview images) or Force Mapping (FM, for force measurements)-mode.

### Generation of *Knock-in* mouse model for Jup S665A mutation

To create the specific mouse strain, where a S665A mutation is introduced into the plakoglobin (JUP) locus, a targeting vector (I143.4 TV, Figure S4A) containing two homology regions of 6,6 kb (long homology arm, LA) and 2,86 kb (short homology arm, SA) framing the neomycin (Neo) resistance cassette was generated. The latter was composed of two FRT sites integrated on both ends of the coding sequence for a Neo-resistant gene expressed under the control of a Simian Virus 40 (SV40) promoter. In order to achieve successful genetic modification of the chromosomal JUP locus, both LA and SA were homologously recombined with the targeted genome sequence in embryonic stem (ES) cells derived from C57B1/6N mice. This event resulted into delivery of heterozygous cells containing both a targeted allele carrying the plakoglobin/JUP mutation, and a wild type (WT) allele. Selected ES cell clones were then injected into blastocysts from grey C57B1/6N mice. The surviving blastocysts were consequently transferred into the uterus of pseudo-pregnant animals, which led to the birth of chimeric heterozygous transgenic mice. The latter were backcrossed with germ-line Flp expressing mice (“Flp deletors”). Upon this breeding scheme, the FRT-flanked Neo was excised by Flp-FRT recombination, initiated by the Flp recombinase delivered from the “Flp deletors”. Thus, black offspring carrying the targeting construct with a deleted Neo cassette were established. These animals were backcrossed to homozygosity for the modified JUP locus.

Detailed description for the generations and validation of the Jup S665A mouse can be found in the Supplementary Materials and Methods. Mice were maintained in breeding facility (Max von Pettenkofer Insitut, LMU Munich) under approved animal breeding proposal (ROB 55.2-2532.Vet_02-19-172).

### Image Processing and Statistical Analysis

Origin (OriginLab, Northhampton, MA, USA) was used for analyzing cAMP Assay and AFM data. For statistics and creating graphs GraphPad Prism (GraphPad Software, San Diego, CA, USA) was used. To compare means of more than 2 samples one-way ANOVA with Bonferroni correction was used and significance was determined for a p-value of 0.5. For two independent groups unpaired T-test was applied. Image processing was performed with Photoshop CS5 (Adobe, San José, USA) and LAS X life science (Leica, Wetzlar, Germany).

## Author contribution

AS, MW, SE, DE, DK, SK, MR and FR conducting experiments and acquiring and analyzing data. MR, KG, ES, AY providing reagents and patients’ material. FV, JW designing research study. FV, JW, AS writing the manuscript. All critically reviewing the manuscript.

AS, MW, SE share first authorship. AS is listed first as she provided her expertise in conducting experiments and acquiring data and mainly analyzed data where she supported MW and SE during their MD thesis.

FV, JW share last authorship. Protective cAMP signaling in pemphigus is a common scientific interest of FV and JW and thus study design and development of the study as well as interpretation of the data and manuscript writing was conducted together. FV and JW benefit from an equal collaboration in terms of different methodical expertise/ focus and supervision of MD students and Post-Docs.

## Acknowledgement

We thank Nadine Albrecht, Martina Hitzenbichler, Sabine Mühlsimer, Kilian Skrowranek and Andrea Wehmeyer for excellent technical assistance. This work was supported by DFG FOR 2497 TP5 to JW and TP6 to FV and the FöFoLe program of the LMU to JW and FV.

## References

1 Green, K. J., Jaiganesh, A. & Broussard, J. A. Desmosomes: Essential contributors to an integrated intercellular junction network. F1000Res 8, doi:10.12688/f1000research.20942.1 (2019).

2 Waschke, J. Desmogleins as signaling hubs regulating cell cohesion and tissue/organ function in skin and heart - EFEM lecture 2018. Ann Anat 226, 96–100, doi:10.1016/j.aanat.2018.11.006 (2019).

3 Broussard, J. A., Getsios, S. & Green, K. J. Desmosome regulation and signaling in disease. Cell Tissue Res 360, 501–512, doi:10.1007/s00441-015-2136-5 (2015).

4 Garrod, D. & Chidgey, M. Desmosome structure, composition and function. Biochim Biophys Acta 1778, 572–587, doi:10.1016/j.bbamem.2007.07.014 (2008).

5 Schmidt, E., Kasperkiewicz, M. & Joly, P. Pemphigus. Lancet 394, 882–894, doi:10.1016/S0140-6736(19)31778-7 (2019).

6 Kasperkiewicz, M. et al. Pemphigus. Nat Rev Dis Primers 3, 17026, doi:10.1038/nrdp.2017.26 (2017).

7 Ellebrecht, C. T. & Payne, A. S. Setting the target for pemphigus vulgaris therapy. JCI Insight 2, e92021, doi:10.1172/jci.insight.92021 (2017).

8 Pollmann, R., Schmidt, T., Eming, R. & Hertl, M. Pemphigus: a Comprehensive Review on Pathogenesis, Clinical Presentation and Novel Therapeutic Approaches. Clin Rev Allergy Immunol 54, 1–25, doi:10.1007/s12016-017-8662-z (2018).

9 Didona, D., Maglie, R., Eming, R. & Hertl, M. Pemphigus: Current and Future Therapeutic Strategies. Front Immunol 10, 1418, doi:10.3389/fimmu.2019.01418 (2019).

10 Heelan, B. T. et al. Effect of anti-CD20 (rituximab) on resistant thrombocytopenia in autoimmune lymphoproliferative syndrome. Br J Haematol 118, 1078–1081 (2002).

11 Ran, N. A. & Payne, A. S. Rituximab therapy in pemphigus and other autoantibody-mediated diseases. F1000Res 6, 83, doi:10.12688/f1000research.9476.1 (2017).

12 Musette, P. & Bouaziz, J. D. B Cell Modulation Strategies in Autoimmune Diseases: New Concepts. Front Immunol 9, 622, doi:10.3389/fimmu.2018.00622 (2018).

13 Joly, P. et al. Updated S2K guidelines on the management of pemphigus vulgaris and foliaceus initiated by the european academy of dermatology and venereology (EADV). J Eur Acad Dermatol Venereol 34, 1900–1913, doi:10.1111/jdv.16752 (2020).

14 Spindler, V. & Waschke, J. Pemphigus-A Disease of Desmosome Dysfunction Caused by Multiple Mechanisms. Front Immunol 9, 136, doi:10.3389/fimmu.2018.00136 (2018).

15 Spindler, V. et al. Mechanisms Causing Loss of Keratinocyte Cohesion in Pemphigus. J Invest Dermatol 138, 32–37, doi:10.1016/j.jid.2017.06.022 (2018).

16 Sajda, T. & Sinha, A. A. Autoantibody Signaling in Pemphigus Vulgaris: Development of an Integrated Model. Front Immunol 9, 692, doi:10.3389/fimmu.2018.00692 (2018).

17 Schmitt, T. & Waschke, J. Autoantibody-Specific Signalling in Pemphigus. Frontiers in Medicine 8, doi:10.3389/fmed.2021.701809 (2021).

18 Schmitt, T. et al. Ca(2+) signalling is critical for autoantibody-induced blistering of human epidermis in pemphigus. Br J Dermatol, doi:10.1111/bjd.20091 (2021).

19 Spindler, V. et al. Peptide-mediated desmoglein 3 crosslinking prevents pemphigus vulgaris autoantibody-induced skin blistering. J Clin Invest 123, 800–811, doi:10.1172/JCI60139 (2013).

20 Egu, D. T., Kugelmann, D. & Waschke, J. Role of PKC and ERK Signaling in Epidermal Blistering and Desmosome Regulation in Pemphigus. Front Immunol 10, 2883, doi:10.3389/fimmu.2019.02883 (2019).

21 Kugelmann, D. et al. Role of Src and Cortactin in Pemphigus Skin Blistering. Front Immunol 10, 626, doi:10.3389/fimmu.2019.00626 (2019).

22 Walter, E. et al. Role of Dsg1- and Dsg3-Mediated Signaling in Pemphigus Autoantibody-Induced Loss of Keratinocyte Cohesion. Front Immunol 10, 1128, doi:10.3389/fimmu.2019.01128 (2019).

23 Burmester, I. A. K. et al. Identification of novel therapeutic targets for blocking acantholysis in pemphigus. Br J Pharmacol 177, 5114–5130, doi:10.1111/bph.15233 (2020).

24 Berkowitz, P. et al. p38MAPK inhibition prevents disease in pemphigus vulgaris mice. Proc Natl Acad Sci U S A 103, 12855–12860, doi:10.1073/pnas.0602973103 (2006).

25 Ivars, M. et al. The involvement of ADAM10 in acantholysis in mucocutaneous pemphigus vulgaris depends on the autoantibody profile of each patient. Br J Dermatol 182, 1194–1204, doi:10.1111/bjd.18382 (2020).

26 Spindler, V., Vielmuth, F., Schmidt, E., Rubenstein, D. S. & Waschke, J. Protective endogenous cyclic adenosine 5’-monophosphate signaling triggered by pemphigus autoantibodies. J Immunol 185, 6831–6838, doi:10.4049/jimmunol.1002675 (2010).

27 Schlegel, N. & Waschke, J. cAMP with other signaling cues converges on Rac1 to stabilize the endothelial barrier-a signaling pathway compromised in inflammation. Cell Tissue Res 355, 587–596, doi:10.1007/s00441-013-1755-y (2014).

28 Schinner, C. et al. Adrenergic Signaling Strengthens Cardiac Myocyte Cohesion. Circ Res 120, 1305–1317, doi:10.1161/CIRCRESAHA.116.309631 (2017).

29 Yeruva, S. et al. Adrenergic Signaling-Induced Ultrastructural Strengthening of Intercalated Discs via Plakoglobin Is Crucial for Positive Adhesiotropy in Murine Cardiomyocytes. Front Physiol 11, 430, doi:10.3389/fphys.2020.00430 (2020).

30 Zebda, R. & Paller, A. S. Phosphodiesterase 4 inhibitors. J Am Acad Dermatol 78, S43–S52, doi:10.1016/j.jaad.2017.11.056 (2018).

31 Li, H., Zuo, J. & Tang, W. Phosphodiesterase-4 Inhibitors for the Treatment of Inflammatory Diseases. Front Pharmacol 9, 1048, doi:10.3389/fphar.2018.01048 (2018).

32 Pincelli, C., Schafer, P. H., French, L. E., Augustin, M. & Krueger, J. G. Mechanisms Underlying the Clinical Effects of Apremilast for Psoriasis. J Drugs Dermatol 17, 835–840 (2018).

33 Torres, T. & Puig, L. Apremilast: A Novel Oral Treatment for Psoriasis and Psoriatic Arthritis. Am J Clin Dermatol 19, 23–32, doi:10.1007/s40257-017-0302-0 (2018).

34 Keating, G. M. Apremilast: A Review in Psoriasis and Psoriatic Arthritis. Drugs 77, 459–472, doi:10.1007/s40265-017-0709-1 (2017).

35 Hatemi, G. et al. Trial of Apremilast for Oral Ulcers in Behcet’s Syndrome. N Engl J Med 381, 1918–1928, doi:10.1056/NEJMoa1816594 (2019).

36 Meier, K., Holstein, J., Solimani, F., Waschke, J. & Ghoreschi, K. Case Report: Apremilast for Therapy-Resistant Pemphigus Vulgaris. Front Immunol 11, 588315, doi:10.3389/fimmu.2020.588315 (2020).

37 Saito, M. et al. Signaling dependent and independent mechanisms in pemphigus vulgaris blister formation. PLoS One 7, e50696, doi:10.1371/journal.pone.0050696 (2012).

38 Shimizu, A. et al. IgG binds to desmoglein 3 in desmosomes and causes a desmosomal split without keratin retraction in a pemphigus mouse model. J Invest Dermatol 122, 1145–1153, doi:10.1111/j.0022-202X.2004.22426.x (2004).

39 van der Wier, G., Pas, H. H., Kramer, D., Diercks, G. F. H. & Jonkman, M. F. Smaller desmosomes are seen in the skin of pemphigus patients with anti-desmoglein 1 antibodies but not in patients with anti-desmoglein 3 antibodies. J Invest Dermatol 134, 2287–2290, doi:10.1038/jid.2014.140 (2014).

40 Sokol, E. et al. Large-Scale Electron Microscopy Maps of Patient Skin and Mucosa Provide Insight into Pathogenesis of Blistering Diseases. J Invest Dermatol 135, 1763–1770, doi:10.1038/jid.2015.109 (2015).

41 Stahley, S. N., Bartle, E. I., Atkinson, C. E., Kowalczyk, A. P. & Mattheyses, A. L. Molecular organization of the desmosome as revealed by direct stochastic optical reconstruction microscopy. J Cell Sci 129, 2897–2904, doi:10.1242/jcs.185785 (2016).

42 Egu, D. T., Walter, E., Spindler, V. & Waschke, J. Inhibition of p38MAPK signaling prevents epidermal blistering and alterations of desmosome structure induced by pemphigus autoantibodies in human epidermis. Br J Dermatol, doi:10.1111/bjd.15721 (2017).

43 Berkowitz, P. et al. Desmosome signaling. Inhibition of p38MAPK prevents pemphigus vulgaris IgG-induced cytoskeleton reorganization. J Biol Chem 280, 23778–23784, doi:10.1074/jbc.M501365200 (2005).

44 Schlogl, E. et al. Keratin Retraction and Desmoglein3 Internalization Independently Contribute to Autoantibody-Induced Cell Dissociation in Pemphigus Vulgaris. Front Immunol 9, 858, doi:10.3389/fimmu.2018.00858 (2018).

45 Vielmuth, F. et al. Keratins Regulate p38MAPK-Dependent Desmoglein Binding Properties in Pemphigus. Front Immunol 9, 528, doi:10.3389/fimmu.2018.00528 (2018).

46 Tsunoda, K. et al. Induction of pemphigus phenotype by a mouse monoclonal antibody against the amino-terminal adhesive interface of desmoglein 3. J Immunol 170, 2170–2178, doi:10.4049/jimmunol.170.4.2170 (2003).

47 Caldelari, R. et al. A central role for the armadillo protein plakoglobin in the autoimmune disease pemphigus vulgaris. J Cell Biol 153, 823–834 (2001).

48 Aoyama, Y. & Kitajima, Y. Pemphigus vulgaris-IgG causes a rapid depletion of desmoglein 3 (Dsg3) from the Triton X-100 soluble pools, leading to the formation of Dsg3-depleted desmosomes in a human squamous carcinoma cell line, DJM-1 cells. J Invest Dermatol 112, 67–71, doi:10.1046/j.1523-1747.1999.00463.x (1999).

49 Vielmuth, F., Waschke, J. & Spindler, V. Loss of Desmoglein Binding Is Not Sufficient for Keratinocyte Dissociation in Pemphigus. J Invest Dermatol 135, 3068–3077, doi:10.1038/jid.2015.324 (2015).

50 Vielmuth, F. et al. Keratins Regulate the Adhesive Properties of Desmosomal Cadherins through Signaling. J Invest Dermatol 138, 121–131, doi:10.1016/j.jid.2017.08.033 (2018).

51 Vielmuth, F., Hartlieb, E., Kugelmann, D., Waschke, J. & Spindler, V. Atomic force microscopy identifies regions of distinct desmoglein 3 adhesive properties on living keratinocytes. Nanomedicine 11, 511–520, doi:10.1016/j.nano.2014.10.006 (2015).

52 Sariisik, E. et al. Decoding Cytoskeleton-Anchored and Non-Anchored Receptors from Single-Cell Adhesion Force Data. Biophys J 109, 1330–1333, doi:10.1016/j.bpj.2015.07.048 (2015).

53 Wilgram, G. F., Caulfield, J. B. & Lever, W. F. An electron microscopic study of acantholysis in pemphigus vulgaris. J Invest Dermatol 36, 373–382 (1961).

54 Radeva, M. Y. & Waschke, J. Mind the gap: mechanisms regulating the endothelial barrier. Acta Physiol (Oxf) 222, doi:10.1111/apha.12860 (2018).

55 Waschke, J. et al. Inhibition of Rho A activity causes pemphigus skin blistering. J Cell Biol 175, 721–727, doi:10.1083/jcb.200605125 (2006).

56 Kroger, C. et al. Keratins control intercellular adhesion involving PKC-alpha-mediated desmoplakin phosphorylation. J Cell Biol 201, 681–692, doi:10.1083/jcb.201208162 (2013).

57 Fischmeister, R. Is cAMP good or bad? Depends on where it’s made. Circ Res 98, 582–584, doi:10.1161/01.RES.0000215564.22445.7e (2006).

58 Spindler, V., Dehner, C., Hubner, S. & Waschke, J. Plakoglobin but not desmoplakin regulates keratinocyte cohesion via modulation of p38MAPK signaling. J Invest Dermatol 134, 1655–1664, doi:10.1038/jid.2014.21 (2014).

59 Muller, E. J., Hunziker, T. & Suter, M. M. Keratin intermediate filament retraction is linked to plakoglobin-dependent signaling in pemphigus vulgaris. J Am Acad Dermatol 56, 890–891; author reply 891-892, doi:10.1016/j.jaad.2006.10.989 (2007).

60 Simard, M. et al. A Tissue-Engineered Human Psoriatic Skin Model to Investigate the Implication of cAMP in Psoriasis: Differential Impacts of Cholera Toxin and Isoproterenol on cAMP Levels of the Epidermis. Int J Mol Sci 21, doi:10.3390/ijms21155215 (2020).

61 Takahashi, H., Kinouchi, M., Tamura, T. & Iizuka, H. Decreased beta 2-adrenergic receptor-mRNA and loricrin-mRNA, and increased involucrin-mRNA transcripts in psoriatic epidermis: analysis by reverse transcription-polymerase chain reaction. Br J Dermatol 134, 1065–1069 (1996).

62 Milakovic, M. & Gooderham, M. J. Phosphodiesterase-4 Inhibition in Psoriasis. Psoriasis (Auckl) 11, 21–29, doi:10.2147/PTT.S303634 (2021).

63 Papp, K. et al. Apremilast, an oral phosphodiesterase 4 (PDE4) inhibitor, in patients with moderate to severe plaque psoriasis: Results of a phase III, randomized, controlled trial (Efficacy and Safety Trial Evaluating the Effects of Apremilast in Psoriasis [ESTEEM] 1). J Am Acad Dermatol 73, 37–49, doi:10.1016/j.jaad.2015.03.049 (2015).

64 Zhang, X. et al. Pemphigus Associated with Psoriasis Vulgaris: A Retrospective Study of Seven Patients and a Review of the Literature. Acta Dermatovenerol Croat 26, 226–232 (2018).

65 Balighi, K., Daneshpazhooh, M., Mahmoudi, H. & Tavakolpour, S. Switching from pemphigus vulgaris to psoriasis: a rare report of three cases. Int J Dermatol, doi:10.1111/ijd.14823 (2020).

66 Kridin, K., Ludwig, R. J., Damiani, G. & Cohen, A. D. Increased Risk of Pemphigus among Patients with Psoriasis: A Large-scale Cohort Study. Acta Derm Venereol 100, adv00293, doi:10.2340/00015555-3607 (2020).

67 Kalantari-Dehaghi, M. et al. Pemphigus vulgaris autoantibody profiling by proteomic technique. PLoS One 8, e57587, doi:10.1371/journal.pone.0057587 (2013).

68 Boukamp, P. et al. Normal keratinization in a spontaneously immortalized aneuploid human keratinocyte cell line. J Cell Biol 106, 761–771, doi:10.1083/jcb.106.3.761 (1988).

69 Sigmund, A. M. et al. Dsg2 Upregulation as a Rescue Mechanism in Pemphigus. Front Immunol 11, 581370, doi:10.3389/fimmu.2020.581370 (2020).

70 Heupel, W. M., Zillikens, D., Drenckhahn, D. & Waschke, J. Pemphigus vulgaris IgG directly inhibit desmoglein 3-mediated transinteraction. J Immunol 181, 1825–1834, doi:10.4049/jimmunol.181.3.1825 (2008).

71 Hartlieb, E. et al. Desmoglein 2 is less important than desmoglein 3 for keratinocyte cohesion. PLoS One 8, e53739, doi:10.1371/journal.pone.0053739 (2013).

72 Renart, J., Reiser, J. & Stark, G. R. Transfer of proteins from gels to diazobenzyloxymethyl-paper and detection with antisera: a method for studying antibody specificity and antigen structure. Proceedings of the National Academy of Sciences 76, 3116–3120, doi:10.1073/pnas.76.7.3116 (1979).

73 Towbin, H., Staehelin, T. & Gordon, J. Electrophoretic transfer of proteins from polyacrylamide gels to nitrocellulose sheets: procedure and some applications. Proc Natl Acad Sci U S A 76, 4350–4354, doi:10.1073/pnas.76.9.4350 (1979).

74 Ebner, A. et al. A new, simple method for linking of antibodies to atomic force microscopy tips. Bioconjug Chem 18, 1176–1184, doi:10.1021/bc070030s (2007).

