## Supplemental Material for "Apremilast prevents blistering in human epidermis by stabilization of keratinocyte adhesion in pemphigus"

### Supplementary Information

### Supplementary Material and Methods

Cell Culture and Test Reagents

HaCaT immortalized human keratinocytes [1] were cultured in Dulbecco's Modified Eagle Medium (Thermo Fisher Scientific, Waltham, Massachusetts) containing 10% FCS (Biochrom, Berlin, Germany), 30 µg/ml penicillin and 65 µg/ml streptomycin (both AppliChem, Darmstadt, Germany) at 37°C in a humidified atmosphere of 5% $CO_2$. For some experiments, HaCaT cells stably transfected with keratin5-YFP (kind gift of Reinhard Windoffer and Nicole Schwarz, Institute of Molecular and Cellular Anatomy, RWTH Aachen University) were used. Normal human keratinocytes (NHEK) were cultured in epidermal keratinocyte medium (CnT-0.7, Cellntec, Bern, Switzerland) containing low calcium (0.06 mM) until 80% confluence and subsequently switched to high $Ca^{2+}$ condition (1.8 mM) for 24 h before experiments were performed.

Murine keratinocytes (MEK) were grown in complete FAD medium (Pan Biotech, Aidenbach, Germany) containing 0.05 mM $Ca^{2+}$ as described in detail before [3]. Experiments were conducted 48 h after cells were switched to high $Ca^{2+}$ FAD-medium ($Ca^{2+}$ concentration: 1.2 mM).

Generation of *Knock-in* mouse model for Jup S665A mutation

To introduce the S665A mutation into the plakoglobin (JUP) locus, a targeting vector (I143.4 TV, Figure S4A) containing two homology regions of 6,6 kb (long homology arm, LA) and 2,86 kb (short homology arm, SA) framing the neomycin resistance cassette was generated. Both, LA and SA, were a product of a PCR where C57Bl/6N BAC DNA was used as a template. Thus, a 2,86 kb NotI - EcoRV fragment spanning exon 9 (E9) and an adjacent 6,6 kb SbfI - AscI fragment containing the genomic region of JUP from exon 10 (E10) to exon 14 (E14) were amplified. Both homologous arms were subcloned on either side of a neomycin (Neo) resistance cassette, introduced previously into pBS-KS vector bone. The Neo cassette was composed of two FRT sites integrated on both ends of the coding sequence for Neo resistant gene expressed under the control of Simian Virus 40 (SV40) promoter. The Neo selection marker was inserted in a less conserved region within the intron 9, in an area about 130 nucleotides upstream of E10. The S665A mutation was introduced within the LA, about 500 bps downstream of the insertion site of the neomycin cassette. More specifically, the mutation is located in exon E11 and it is caused by a transition of the first T to G in the TCT codon, encoding for serine (Ser) in CDS position 1993. This resulted in generation of GCT nucleotide triplet which codes for alanine (Ala) (Figure S4A). To incorporate the mutation, an overlay-extension PCR was performed. Hence, two PCRs were run in parallel using different primer pairs (pair 1: I143.4/I143.6 and pair 2: I143.5/I143.7; Table 1). The resulting PCR products served as a template for a third PCR with the primers set I143.4/I143.5 (Table 1 and Figure S4B). A PCR fragment, spanning E10 and mutation-carrying E11 was amplified and subsequently cloned into the LA, within the I143.4 TV targeting vector, by using SbfI and BstEII restriction sites. As a final step, to verify the integrity of the vector bone and the newly inserted regions, in addition to the restriction analysis, a complete sequencing of the key regions within the targeting vector was done.

Detection of homologous recombination in embryonic stem cells

C57BL/6N-derived embryonic stem (ES) cells were transfected with I143.4 TV targeting vector. This generated heterozygous cells containing both a targeted allele carrying the plakoglobin/JUP mutation, and a wild type (WT) allele (Figure S4B). Prior to the electroporation of ES cells, the targeting vector was linearized with the NotI restriction enzyme. For selection and maintenance of the ES cells transfected with the vector carrying a Neo resistance gene, G418 was added at a final concentration of 0,2 mg/ml. Following 8 days of selection, a total of 192 clones were picked and tested to verify a successful homologous recombination. For this purpose, a screening PCR was established, where a newly constructed primer pair (I143.9/518LRPCR2) was designed to align only to the targeted allele, but not to the WT allele (Table 1 and Figure S4B). Additionally, the primer set was selected so that it gives an amplicon only when a recombination event occurs in the targeted locus, but not in any other random locus. Therefore, the 518LRPCR2 primer specifically binds to the vector and not to the genome; within a region of the neomycin cassette. In contrast, the I143.9 primer aligns to the genome only, more precisely, to a sequence that is not part of the targeting vector. Thus, if a signal was detected, an association between the vector and the Jup locus could be validated. To test the specificity of the primers, a control vector (I143.2 CV) was generated, which was later used as an internal positive control for the screening PCR. The vector contains an elongated SA homology region, which holds 50 bp of extra genomic sequence that is not present in the I143.4 TV targeting vector and is used as a template to which a I143.9 primer is selectively aligning. In addition to the screening PCR analysis, the homologous recombination in the ES was confirmed by sequencing and Southern blot analysis.

Southern blot analysis

The validation of positive clones carrying correct homologous recombination with the Jup locus was achieved by Southern blot analysis.

DNA from individual clones was digested with the restriction enzyme NsiI, the location of the sites are shown in Figure S4B, confirming that, compared to the WT, the targeted allele (TG) has additional NsiI restriction sites, specifically localized within the neomycin cassette. NsiI-digested genomic DNA was then analyzed using [α-32P] dCTP DNA 3’ probe that corresponds to the genomic sequence downstream of the incorporated mutation (Figure S4B). The 3’ external probe (516 bp in length) was generated by PCR using the primer combination I143.21/I143.22 (Table 1) and genomic DNA as a template. The probe was located approximately 3,4 kb downstream of E14 and about 880 bp downstream from the end of LA homologous arm. After gel extraction and purification, the probe was used to identify the complementary DNA fragment of interest. Due to the size of the Neo cassette, the TG allele is larger than the WT allele, therefore, the size of the restriction fragment detected by the probes was 9,9 kb. In contrast, presence of a signal corresponding to 8.6 kb, confirmed the existence of a WT allele.

Detection of S665A mutation in the ES cell genome

To test for the presence of the mutation in ES cells, a region surrounding the mutation was amplified using a forward primer (I143.4), specific for region in intron 9, and a reverse primer (I143.5), binding to intron 11 (Table 1 and Figure S4B). Designed primers yielded a PCR amplification product with a size of 1094 bp in both, TG- and the WT- allele. The 1094-bp sized fragment was purified from a gel and subsequently sequenced with either of the primers used for its amplification.

Blastocyst injection and breeding

Selected ES cell clones were injected into blastocysts from grey C57BL/6N mice. The surviving blastocysts were then transferred into the uterus of pseudo-pregnant animals (3 CD-1 foster mice). This resulted in the birth of chimeric heterozygous transgenic mice, which were then backcrossed with germ-line Flp expressing mice ("Flp deletors"). Upon this breeding scheme, the FRT-flanked Neo was excised by Flp-FRT recombination, initiated by the Flp recombinase delivered from the "Flp deletors". This led to the establishment of black offspring carrying the targeting construct with a deleted Neo cassette.

Genotyping of constitutive JUP Ser665Ala KI mutant

For genotyping of pups from the F1 and F2 generation, genomic DNA extracted from tail or ear tissue was use as a template for a series of different PCRs.

By using the I143.27/I143.28 primer combination (Table 1 and Figure S4C), the samples were screened for the Flp-mediated deletion of the Neo cassette, which is shown by the presence of the remaining FRT site. Positive animals are germline transmitters of the targeted Jup allele after deletion of the Neo cassette. The predicted amplicon for WT allele is 568 bp in size, while for the targeted deleted Neo (TG del-neo), a 686 bp fragment is expected.

To identify offspring that still contained the Neo cassette, the primer combination Neo.MP1/Neo.MP5/Neo.MP6 (Table 1 and Figure S4C) was used for screening. In contrast to Neo.MP1, which binds to a sequence from the Neo cassette and to a region within chromosome 3, Neo.MP5 and Neo.MP6 are unique and align to either of the above mentioned regions. Therefore, whereas both primers of the set Neo.MP1/Neo.MP6 bind to regions within the Neo cassette (expected product of 512 bp), the primer combination Neo.MP1/ Neo.MP5 align to regions within the chromosome 3 (PCR product with expected size of 380 bp, used as an internal control). Additionally, the mice were screened for the presence of the Flp recombinase using the primer combination SD24/SD25, which results in a 568 bp-sized PCR fragment. Finally, to test for the presence of the S665A mutation in Jup Ex 11, amplification of a region (875 bp in size) spanning the mutation was screened by using I143.25/I143.26 primer set (Table 1 and Figure S4C).

Table 1: List of used primer

| Purpose/Aim | Primer name | Sequence 5’-> 3’ |
|---|---|---|
| To generate the short arm (SA) | I143.1 | CAGGATATCGCCCCTAATACCTGAGC |
| To generate the short arm (SA) | I143.2 | CAAGCGGCCGCCTCCTGACTGCTGAGAATAC |
| To generate the long arm (LA) | I143.8 | CTAGGCGCGCCGGACTCTTGGCTGTAGTAG |
| To test the presence of mutation in ES cells<br>To generate the long arm (LA)<br>Overlay-extension PCR | I143.4 | GTTCCTGCAGGGTGTCTGCTCAGGTATTAGG |
| To test the presence of mutation in ES cells<br>Overlay-extension PCR | I143.5 | CAAGGCGCGCCAGAGGTCCTGAGTTCAAGTC |
| Generation of probe for Southern Blot | I143.21 | AGGCTGTCGAGATGATTC |
| Generation of probe for Southern Blot | I143.22 | CAGAGATCAGCCTGTCTATG |
| Overlay-extension PCR | I143.6 | GAATTGGTGAGCTCCACAGCCACTCGCTTACGGTAATC |
| Overlay-extension PCR | I143.7 | GATTACCGTAAGCGAGTGGCTGTGGAGCTCACCAATTC |
| Screening for Neo resistance cassette deletion | I143.27 | GACACATGCGAACATACC |
| Screening for Neo resistance cassette deletion | I143.28 | CCTGTCTAGCCATCTTAGG |
| Screening for Neo resistance cassette | Neo.MP1 | GCTGTGCTCCACGTTGTCAC |
| Screening for Neo resistance cassette | Neo.MP5 | GGAAAGCTGGGCTTGCATCTC |
| Screening for Neo resistance cassette | Neo.MP6 | GGAGCGGCGATACCGTAAAG |
| Screening for Flp recombinase allele | SD24 | CTAATGTTGTGGGAAATTGGAGC |
| Screening for Flp recombinase allele | SD25 | CTCGAGGATAACTTGTTTATTGC |
| Detection of the S665A mutation in mice | I143.25 | CTAGCTCCTGTACTCCTCTG |
| Detection of the S665A mutation in mice | I143.26 | TGGTGGCTCATGACTCTC |
| Screening PCR | I143.9 | GTAACCCAGGCTGTCCTTAG |
| Screening PCR | 5182LRPCR2 | GTTGTGCCCAGTCATAGCCGAATAG |
| presence of Flp recombinase allele | SD24 | CTAATGTTGTGGGAAATTGGAGC |
| presence of Flp recombinase allele | SD25 | CTCGAGGATAACTTGTTTATTGC |

Purification of patients IgG fractions and AK23

We received sera samples of healthy volunteers and Pemphigus patients, who gave their written informed consent for research use (Enno Schmidt, Department of Dermatology, University of Lübeck). The phenotype of the disease was clinically and histopathologically characterized and the antibody profile was measured using Dsg1 and Dsg3-ELISA (Euroimmun, Lübeck, Germany). Titers of PV-IgGs used in this study are: PV1-IgG: 760 U/ml (Dsg1) and 4711 U/ml (Dsg3), PV2-IgG: 1207 U/ml (Dsg1) and 3906 U/ml (PV-IgG2). We purified the IgG-fraction using protein A affinity chromatography. Serum was incubated with protein A agarose (Merck, Darmstadt, Germany) in purification columns for two hours, washed with PBS, eluted with 20 mmol/l sodium citrate solution, neutralized in 2 mol/l sodium carbonate solution and concentrated in a filter unit (Amicon® Ultra-4, Merck, Darmstadt, Germany) at 19000 g for 20 min. Afterwards the antibodies were buffered in PBS and stored at -20°C. To test their functionality, we performed a Dispase II based dissociation assay.

AK23, a pathogenic monoclonal anti-Dsg-3-Antibody produced in a mouse model, was bought from Medical & Biological Laboratories Co., Ltd. (Nagoya, Japan).

Purification of recombinant Dsg-Fc constructs

Purification of recombinant Dsg-Fc proteins was carried out as described before [4]. Briefly, Dsg3- extracellular domain-Fc constructs were expressed stably in Chinese hamster ovarian cells (CHO cells). After reaching 65-75 % confluence, supernatant was collected and recombinant proteins were isolated by protein A agarose affinity chromatography (Life Technologies/ThermoFisher scientific, Waltham, USA; details please refer purification of Antibodies). Coomassie staining and Western blot analysis with aDsg3 mAb (clone5G11; Life Technologies) were performed to test purity.

*Ex-vivo* pemphigus skin model

Human *ex vivo* pemphigus model was performed with cadavers of the human body donor program without history of skin diseases from the Institute of Anatomy and Cell Biology, Ludwig-Maximilians-University Munich, Germany as described before [5]. Skin samples were taken from body donors who had died no longer than 24 hours before and had no history of skin lesions. A 30.5 G needle (B.Braun, Melsungen, Germany) was used to inject either 50 µl Apremilast in DMSO or DMSO and after 1 h of pre-incubation another 50 µl of either PV- or C-IgG. Samples were incubated in Dulbecco's modified Eagle's medium (DMEM) in a humidified atmosphere of 95% air and 5% $CO_2$ at 37°C for 24 h. After incubation a defined shear stress was applied using a rubber head. Subsequently, specimen were either embedded in Tissue-tec (Leica, Wetzlar, Germany) for cryosectioning or in glutaraldehyde for electron microscopy analysis.

Electron microscopy

Electron microscopy analyses

The treated skin pieces, 24 h after incubation, were dissected into small pieces (approximately 2 mm) and fixed by immersion in 2.5% glutaraldehyde in PBS for 1 h at room temperature. The samples were either transferred to PBS for storage at 40°C or subsequently processed as described in Egu et al., 2017 [5]. Briefly, they were taken through three washes of PBS buffer and post-fixed in 2% osmium tetroxide for 3 h. This was followed by successive washings in ascending ethanol series and finally cleared in propylene oxide. The samples were finally embedded in EPON 812 resin (SERVA Electrophoresis GmBH, Heidelberg, Germany), and cured as per the company’s recommendation. The resulting blocks were sliced at 60 nm thickness using ultramicrotome (Reichert-Jung Ultracut E, Optische Werke AG, Vienna, Austria) with a diamond knife (DiATOME Electron Microscopy Sciences, Hatfield, PA). Silver shining sections were harvested using 150 mesh copper/rhodium grids (Plano GmbH, Wetzlar, Germany). Finally, sections were contrasted using uranyl acetate and lead citrate. Images were captured at 4000x and 12,500x magnification with a Libra 120 transmission electron microscopy (Carl Zeiss NTS GmbH, Germany) equipped with a SSCCD camera system (TRS, Olympus, Tokyo, Japan).

Ultrastructural quantification

Electromicrographs were taken along the cleavage plane of pemphigus vulgaris in the deep layer of the epidermis. Accordingly, desmosomes between adjacent basal keratinocytes as well as those between basal and suprabasal keratinocytes were considered in the evaluation. Images were captured at 4000x magnification, and for each set of experiment 20 -30 images and ~100 desmosomes were evaluated per condition using ImageJ software (Wayne Rasband; https://imagej.nih.gov/ij/). Measurement of desmosome size and number as well as quantification of split desmosomes and retracted keratin filaments in the vicinity of blister formation was performed as previously described [6].

Cell lysis and Western blotting

For protein analysis cells were washed and lysed in 60 µl SDS-lysis-buffer (25 mM HEPES, 2 mMol EDTA, 25 mM NaF and 1 % sodiumdodecylsulfate, pH 7.6, protease inhibitors). For some experiments cells were incubated with extraction buffer (0.5% Triton-X-100, 50 mM MES , 25 mM EGTA, 5 mM $MgCl_2$, pH = 6.8, protease inhibitors Aprotenin, Pepstatin, Leupeptin, and phenylmethylsulfonylfluoride) for 20 minutes on ice, scraped and afterwards centrifugated at 19.000 g to split triton-soluble non-cytoskeletal bound from the cytoskeletal bound non-soluble proteins. Supernatants containing the Triton-soluble fraction were collected and pellet was resuspended in SDS-lysis-buffer.

For detection of phosphorylated proteins cells were lysed in laemmli-buffer containing 0.5% Triton-X-100, 50 mM MES , 25 mM EGTA, 5 mM $MgCl_2$, pH = 6.8, protease inhibitors Aprotenin, Pepstatin, Leupeptin, and phenylmethylsulfonylfluoride, phosphatase inhibitor mix (1 Tbl. per 10 ml buffer, Phosstop Phosphatase Inhibitor Cocktail Tablets, Roche, Mannheim) and 50 mM dithiothreitol, sonicated and heated at 95° C for 5 min. Afterwards a special Manganese (II)-Phos-tagTM SDS-Page (FUJIFILM Wako Chemicals Europe GmbH, Neuss) was used which creates phospho-binding sites by addition of phos-tag acrylamide and manganese metal ions. Thus, phosphorylated proteins run more slowly in the gel than the unphosphorylated form. A gel containing water instead of phos-tag-acrylamid was used for control.

Electrophoresis and western blot were performed as standard protocols as described elsewhere [7,8]. Nitrocellulose-membranes were used (Thermo Fisher Scientific, Waltham, Massachusetts). The following primary antibodies were used: anti-Dsg3 rabbit polyclonal (Biozol, Eching, Germany), anti-GAPDH mouse (Santa Cruz, Dallas, TX), anti-plakoglobin mouse (Progen, Heidelberg, Germany), anti-desmoplakin rabbit polyclonal (Abclonal, MA, USA), anti-α-Tubulin and anti-CK14 monoclonal (Abcam, Berlin, Germany), β-catenin monoclonal (BD, Eysins, Switzerland) anti-p-Plakoglobin S665 [9]. After washing the following HRP-coupled secondary antibodies have been used: peroxidase-coupled-goat-anti-rabbit or –goat-anti-mouse (Dianova, Hamburg, Germany) Chemiluminescence was detected using the ECL-system.

<u>Immunostaining</u>

For immunostaining cells were fixed with PFA 2% for 10 min, permeabilized with 0.1% Triton-X-100 for 5 min and treated with bovine serum albumin and normal goat serum to block unspecific antibody binding. Additionally, tissues were fixed in PFA 2% for 15 min, permeabilized with 1% Triton-X-100 for 40 min and treated with bovine serum albumin in combination with normal goat serum to block unspecific antibody binding.

Subsequently, cells or tissues were incubated at 4° with the following primary antibodies: anti-desmoglein-3 rabbit polyclonal (Biozol, Eching, Germany), anti-desmoglein-3 mouse monoclonal (Invitrogen, Carlsbad, CA), anti-desmoglein-1 rabbit polyclonal (Abclonal, MA, USA), anti-desmoglein-1 (Progen, Heidelberg, Germany), anti-desmoglein-2 mouse monoclonal (Origene, Rockville, MD), anti-E-cadherin mouse monoclonal (BD, Eysins,

Switzerland), β-catenin mouse monoclonal (BD, Eysins, Switzerland), anti-panCK-FITC mouse monoclonal (Sigma Aldrich, St. Louis, MO), anti-panCK mouse monoclonal (Sigma Aldrich, St. Louis, MO), anti-CK14 mouse monoclonal (Abcam, Berlin, Germany), Phalloidin Alexa 488 (Molecular Probes, Eugene, OR) anti-plakoglobin mouse monoclonal (Progen, Heidelberg, Germany), anti-p-plakoglobin S665 [9] and anti-Vinculin rabbit polyclonal (Sigma, Aldrich, St. Louis, MO). As secondary antibodies 488-, Cy2- or Cy3 coupled goat anti rabbit or goat anti mouse secondary antibodies (Dianova, Hamburg, Germany) (all dianova, Hamburg, Germany) were applied. 4′,6-Diamidin-2-phenylindol was added to secondary antibody incubation (DAPI, 1 mg/ml) for 10 min. Afterwards probes were mounted with NPG (1% n-Propylgallat und 60% Glycerin in PBS). Images were acquired with a Leica SP5 confocal microscope equipped with a 63× oil objective (Leica, Wetzlar, Germany).

AFM measurements

AFM measurements on living HaCaT keratinocytes were performed as described before in detail [10,11]. Briefly, a NanoWizard® 3 AFM (JPK-Instruments, Berlin, Germany) mounted on an inverted optical microscope (Carl Zeiss, Jena, Germany) or Nanowizard 4 AFM (JPK-Instruments, Berlin, Germany) mounted on an inverted optical microscope (IX73 Olympus, Hamburg, Germany), which both allow selection of the scanning area by visualizing the cells with a 63x objective were used. Pyramidal-shaped D-Tips of Si3N4 MLCT cantilevers (Bruker, Mannheim, Germany) with a nominal spring constant of 0.03 N/m and tip radius of 20 nm were functionalized with purified Dsg3-Fc (concentration of 0.15 mg/ml) as described before [12]. Here, a flexible heterobifunctional acetal-polyethylenglycol (PEG, BroadPharm, San Diego, US) was interspaced between the tip and the recombinant Dsg3-Fc.

AFM was used performed either in Quantitative Imaging (QI, for overview images) or Force Mapping (FM, for force measurements)-mode. In QI mode an area of 20 x 30 µm was sampled with setpoint of 0.3 nN and a pulling speed of 50 µm/s; a Z-length of 2 µm and an additional retract of 150 nm. These settings were applied to avoid cell alterations due to mechanical stress and allow to identify areas of cell-cell-contact further referred as cell borders. In FM mode each map consists of distinct force-distance cycles with each cycle representing one pixel of the map covering an area of 1.25 x 5 µm along a cell border. Setting were adapted to setpoint 0.5 nN, pulling speed 10 µm/s, Z-length 2µm and a resting contact time 0.1 s.

References


1 Boukamp, P. *et al.* Normal keratinization in a spontaneously immortalized aneuploid human keratinocyte cell line. *J Cell Biol* **106**, 761-771, doi:10.1083/jcb.106.3.761 (1988).

2 Walter, E. *et al.* Role of Dsg1- and Dsg3-Mediated Signaling in Pemphigus Autoantibody-Induced Loss of Keratinocyte Cohesion. *Front Immunol* **10**, 1128, doi:10.3389/fimmu.2019.01128 (2019).

3 Seltmann, K. *et al.* Keratins Mediate Localization of Hemidesmosomes and Repress Cell Motility. *Journal of Investigative Dermatology* **133**, 181-190, doi:10.1038/jid.2012.256 (2013).

4 Heupel, W. M., Zillikens, D., Drenckhahn, D. & Waschke, J. Pemphigus vulgaris IgG directly inhibit desmoglein 3-mediated transinteraction. *J Immunol* **181**, 1825-1834, doi:10.4049/jimmunol.181.3.1825 (2008).

5 Egu, D. T., Walter, E., Spindler, V. & Waschke, J. Inhibition of p38MAPK signalling prevents epidermal blistering and alterations of desmosome structure induced by pemphigus autoantibodies in human epidermis. *Br J Dermatol* **177**, 1612-1618, doi:10.1111/bjd.15721 (2017).

6 Egu, D. T. *et al.* A new ex vivo human oral mucosa model reveals that p38MAPK inhibition is not effective in preventing autoantibody-induced mucosal blistering in pemphigus. *Br J Dermatol* **182**, 987-994, doi:10.1111/bjd.18237 (2020).

7 Renart, J., Reiser, J. & Stark, G. R. Transfer of proteins from gels to diazobenzyloxymethyl-paper and detection with antisera: a method for studying antibody specificity and antigen structure. *Proc Natl Acad Sci U S A* **76**, 3116-3120, doi:10.1073/pnas.76.7.3116 (1979).

8 Towbin, H., Staehelin, T. & Gordon, J. Electrophoretic transfer of proteins from polyacrylamide gels to nitrocellulose sheets: procedure and some applications. *Proc Natl Acad Sci U S A* **76**, 4350-4354, doi:10.1073/pnas.76.9.4350 (1979).

9 Yeruva, S. *et al.* Adrenergic Signaling-Induced Ultrastructural Strengthening of Intercalated Discs via Plakoglobin Is Crucial for Positive Adhesiotropy in Murine Cardiomyocytes. *Front Physiol* **11**, 430, doi:10.3389/fphys.2020.00430 (2020).

10 Vielmuth, F., Waschke, J. & Spindler, V. Loss of Desmoglein Binding Is Not Sufficient for Keratinocyte Dissociation in Pemphigus. *J Invest Dermatol* **135**, 3068-3077, doi:10.1038/jid.2015.324 (2015).

11 Vielmuth, F., Hartlieb, E., Kugelmann, D., Waschke, J. & Spindler, V. Atomic force microscopy identifies regions of distinct desmoglein 3 adhesive properties on living keratinocytes. *Nanomedicine* **11**, 511-520, doi:10.1016/j.nano.2014.10.006 (2015).

12 Ebner, A. *et al.* A new, simple method for linking of antibodies to atomic force microscopy tips. *Bioconjug Chem* **18**, 1176-1184, doi:10.1021/bc070030s (2007).

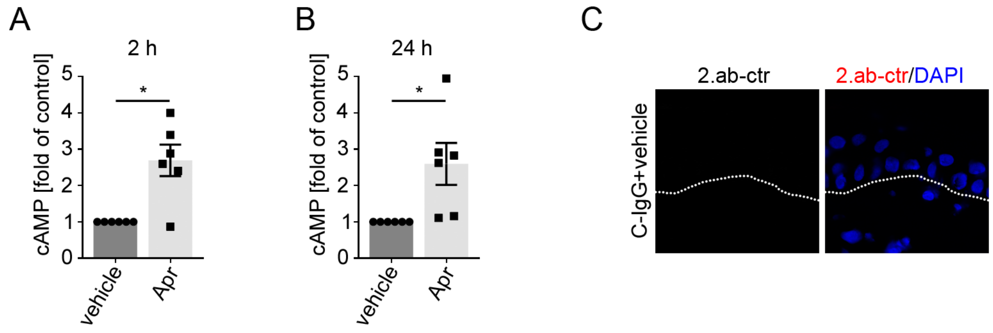


**Figure S1: cAMP ELISA and secondary antibody controls.**

cAMP ELISA of HaCaT cell lysates after incubation with apremilast or vehicle (DMSO) for 2 h (**A)** or 24 h **(B)** reveals a significant increase of cAMP levels after apremilast treatment under both conditions (n=6). Columns indicate mean value normalized to control ± SEM. *P<0.05. Students' T-test. **(C)** Secondary antibody control of immunofluorescence of *ex vivo* skin samples. Representative of n = 4. White dotted line marks basement membrane.

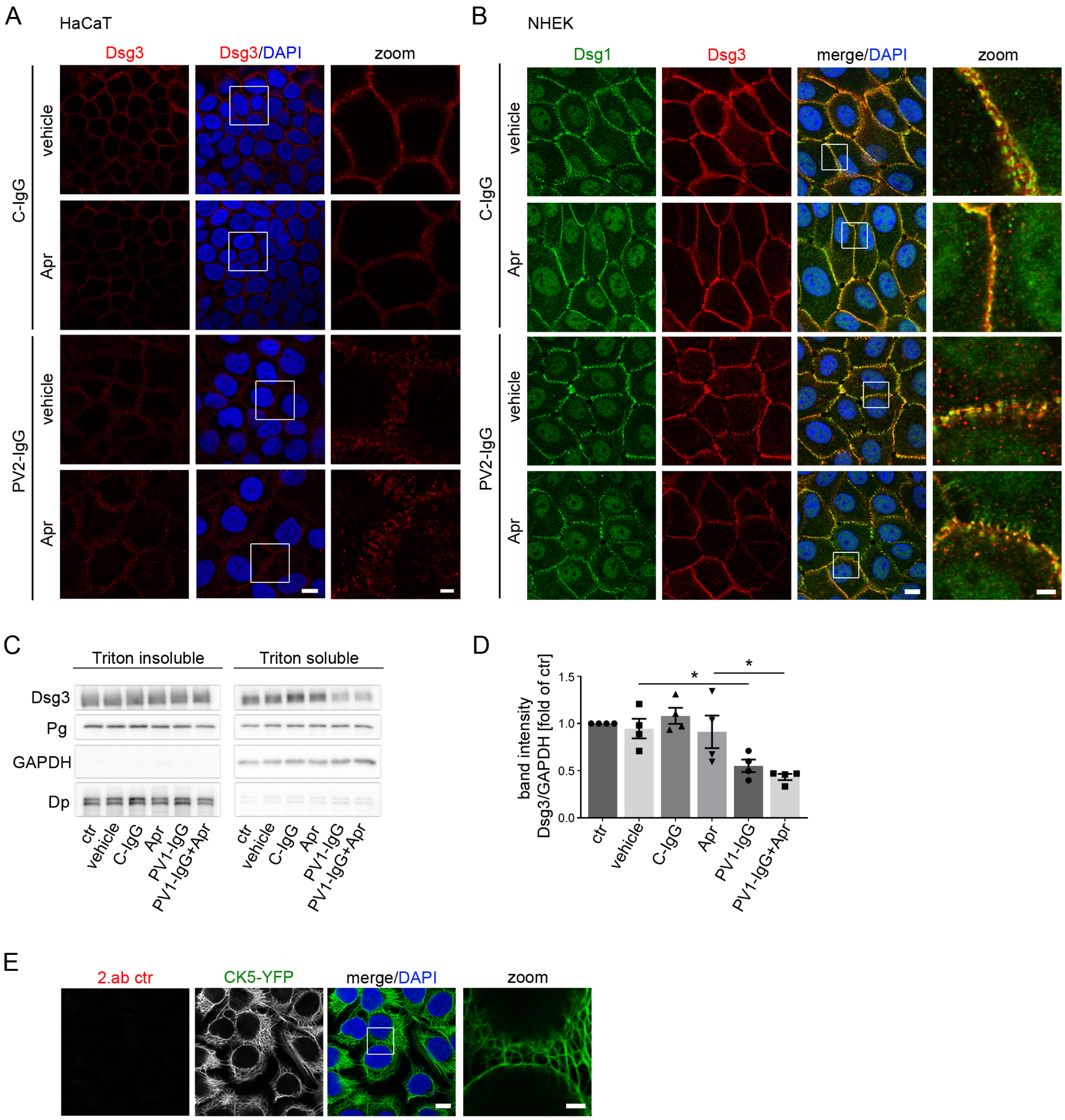


**Figure S2: Apremilast does not rescue PV-IgG-induced fragmentation of Dsg1 and Dsg3 stainings.**

Immunostaining against Dsg3 in HaCaTs **(A)** or Dsg1 and Dsg3 in NHEKs **(B)**. PV2-IgG incubation for 24 h leads to fragmentation of Dsg1 and Dsg3 stainings. 1 h pre-incubation of apremilast does neither ameliorate PV2-IgG induced fragmentation of Dsg1 nor Dsg3 staining. Scale bar = 10 µm. White rectangles depict areas for zoom in. Scale bar (zoom) = 3 µm (HaCaT)/ 2.5 µm (NHEK). Representative of n = 4. **(C)** Representative Western blot of HaCaT Triton soluble and Triton insoluble, desmosome containing fractions shows no effect of apremilast on PV1-IgG-induced Dsg3 depletion. Desmoplakin (Dp) and Glycerol-3-phosphate dehydrogenase (GAPDH) served as loading controls for the respective fraction. **(D)** Quantification of Triton soluble fraction of Western blots (n=4). **(E)** Secondary antibody control of immunofluorescence of HaCaTs stable expressing CK5-YFP. Representative of n = 4. Scale bar = 10 µm. Areas for zoom in marked with white rectangles. Scale bar (zoom) = 2.5 µm. Columns indicate mean values normalized to control ± SEM. *P<0.05. One-way ANOVA with Bonferroni correction. plakoglobin (Pg)

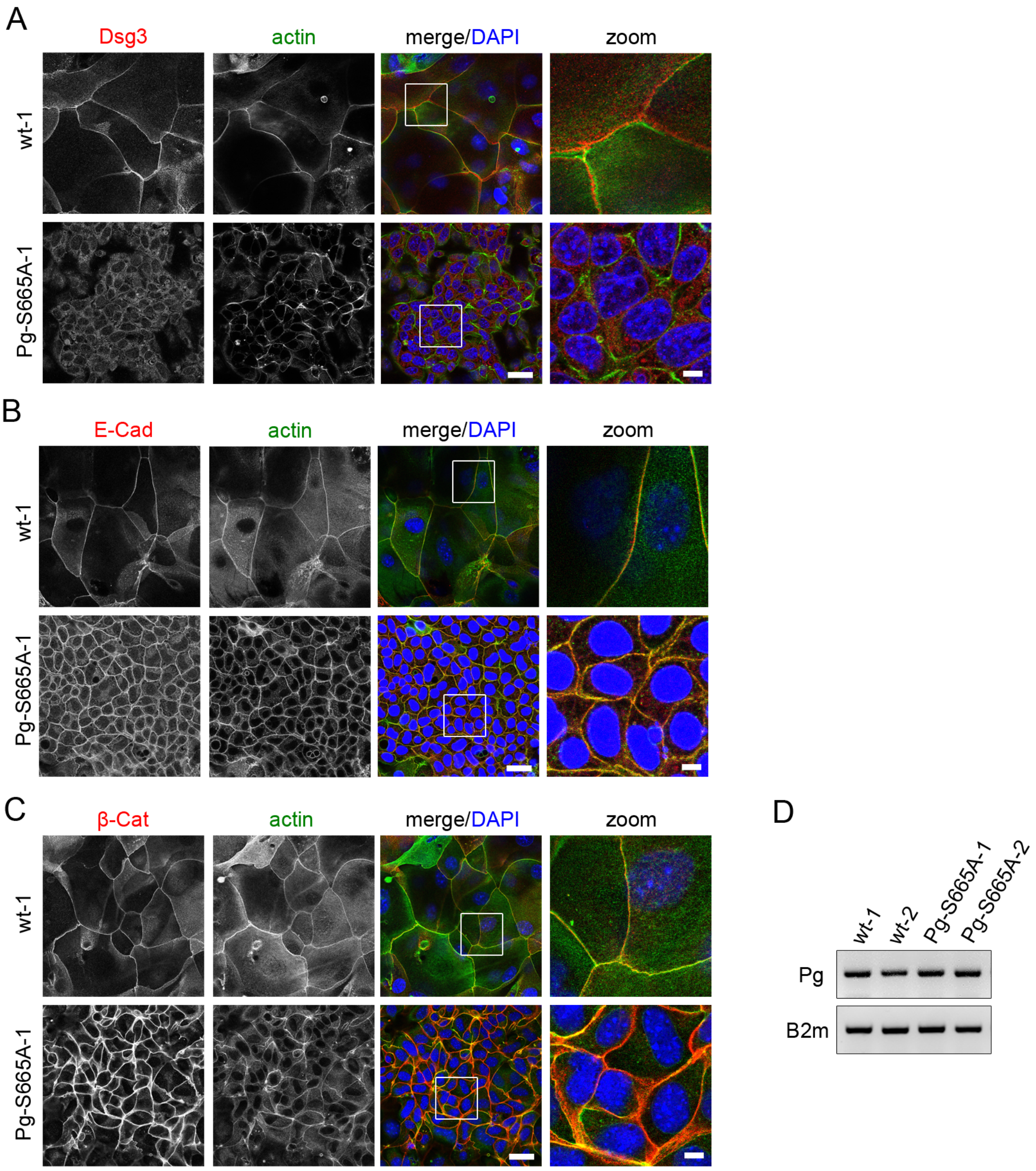


**Figure S3: Keratinocytes phospho-deficient at Pg S665 show alterations in desmosomal protein distribution and compensatory effects on adherens junction proteins.**

**(A)** Co-staining of Dsg3 and actin reveals impaired and in part fragmented distribution of Dsg3 along the cell borders in Pg-S665A keratinocytes. **(B)** Co-staining of E-cadherin (E-Cad) and actin shows a slight upregulation of E-cadherin in Pg-S665A keratinocytes. **(C)** Similarly, co-staining of β-catenin and actin demonstrates a clear upregulation of β-catenin along cell borders in Pg-S665A keratinocytes. **(A-C)** The actin cytoskeleton is broadened along cell borders in Pg-S665A keratinocytes. Scale bar = 25 µm. White rectangles depict areas chosen for zoom ins. Scale bar (zoom) = 5 µm. Representatives of n=4. **(D)** mRNA of *Pg* is unaltered in Pg-S665A keratinocytes compared to wt. *Beta-2-microglobulin (B2m)* was used as in put control. Representative of n=3-4.

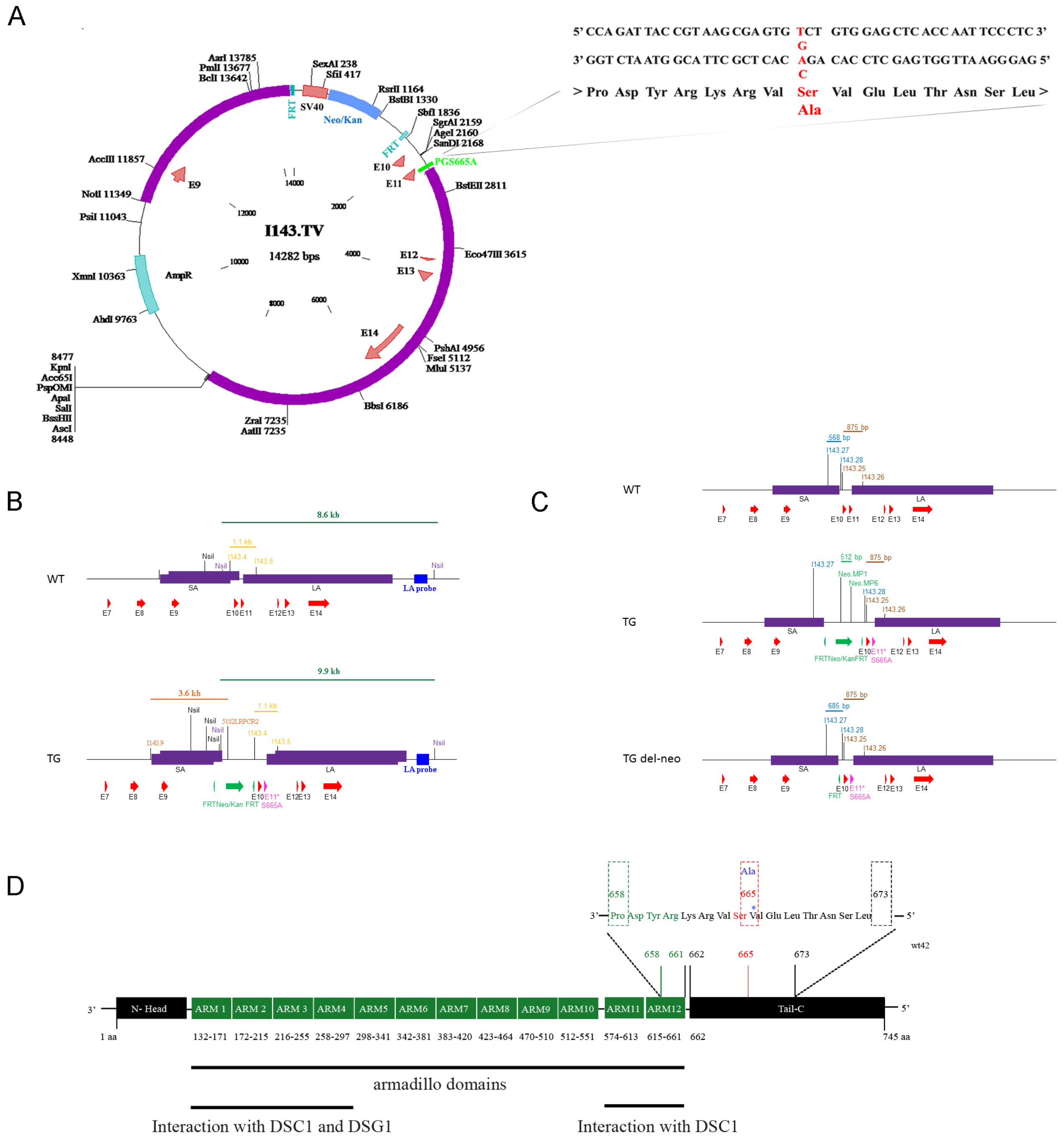

### Figure S4: Generation of Knock-in mouse model for Jup S665A mutation

**(A)** The mutated Jup allele. The 1993 T>G mutation in exon 11 results in the exchange of serine (Ser) for alanine (Ala, red. **(B)** Schematic drawing of the different Jup alleles. The upper panel shows the wild type (WT) allele, the lower panel shows the targeted (TG) allele. Homologous regions (SA: short arm of homology; LA: long arm of homology) used for recombination are depicted as purple boxes. The mutation in exon 11 (E11* S665A, pink) is inserted via homologous recombination together with a FRT-flanked neomycin cassette (green). The primers I143.9 and 5182LRPCR2 were used for PCR screening of recombined ES cells. The corresponding fragment size is indicated in orange. Primers used for confirmation of the S665A mutation are indicated in yellow. The restriction enzyme NsiI, used for Southern blot analysis, and the corresponding 3' external probe (LA probe) and fragments are indicated in blue. **(C)** Schematic drawing of the WT and the targeted locus of plakoglobin, with (TG) and without the neomycin resistance cassette (TG del-neo). The primers used for genotyping and the corresponding amplicon sizes (blue: del-neo PCR, brown: PCR for confirmation of the S665A mutation, green: neomycin PCR) are mapped on the different loci. **(D)** Introduction of S665A mutation into the plakoglobin (JUP) locus. The phospho-site of interest is located at S665 in the C-terminal tail region.
